# The Chloroplast RNA Binding Protein CP29A supports *rbcL* expression during cold acclimation

**DOI:** 10.1101/2023.11.24.568587

**Authors:** Benjamin Lenzen, Florian Rösch, Hannes Ruwe, Nitin Kachariya, Julia Legen, Michael Sattler, Ian Small, Christian Schmitz-Linneweber

## Abstract

The chloroplast genome encodes key components of the photosynthetic light reaction machinery as well as the large subunit of the enzyme central for carbon fixation, RuBisCo. Its expression is predominantly regulated post-transcriptionally, with nuclear-encoded RNA binding proteins (RBPs) playing a key role. Mutants of chloroplast gene expression factors often exhibit impaired chloroplast biogenesis, especially in cold conditions. Low temperatures pose a challenge for plants as this leads to electron imbalances and oxidative damage. A well-known response of plants to this problem is to increase the production of RuBisCo and other Calvin Cycle enzymes in the cold, but how this is achieved is unclear. The chloroplast RBP CP29A has been shown to be essential for cold resistance in growing leaf tissue of *Arabidopsis thaliana*.

Here, we examined CP29A-RNA interaction sites at nucleotide resolution. We discovered that CP29A preferentially binds to the 5’-UTR of *rbcL*, downstream of the binding site of the pentatricopeptide repeat (PPR) protein MRL1. MRL1 is an RBP known to be necessary for the accumulation of *rbcL*. In *Arabidopsis* mutants lacking CP29A, we were unable to observe significant effects on *rbcL*, possibly due to CP29A’s restricted role in a limited number of cells at the base of leaves. In contrast, CRISPR/Cas9-induced mutants of tobacco NtCP29A exhibit cold-dependent photosynthetic deficiencies throughout the entire leaf blade. This is associated with a parallel reduction in *rbcL* mRNA and RbcL protein accumulation. Our work unravels the molecular player behind cold acclimation of the photosynthetic dark reaction.

**Significance Statement:** This study unveils the critical role of CP29A, a chloroplast-localized RNA binding protein, in facilitating plants’ acclimation to cold environments. Through advanced molecular techniques, we discovered that CP29A specifically targets the rbcL mRNA, vital for the production of RuBisCo—a key enzyme in photosynthesis and the most abundant protein on Earth. Our findings elucidate a previously unknown mechanism of how plants adjust to cold stress by regulating RuBisCo levels, highlighting the intricate interplay between nuclear and chloroplast genomes. This research not only advances our understanding of plant cold acclimation but also provides insights that could help enhance plant resilience and productivity when facing temperature challenges.

## Introduction

The chloroplast is crucial in how plants adapt to cold stress (1). Cold tolerance in many plants relies on photosynthetic acclimation during low-temperature growth. When exposed to light in cold conditions, the Calvin cycle enzyme activity declines and thus the availability of electron acceptors becomes limited, while at the same time the plastoquinone pool is fully reduced. This leads to a buildup of reactive oxygen species, triggering signals that control nuclear gene expression. The chloroplast’s response to cold directly affects a plant’s ability to withstand low temperatures (1), indicating its adjustment is crucial for overall resilience to cold.

Chloroplasts undergo intricate adjustments in gene expression to acclimate to cold temperatures, with critical roles played by transcripts encoding key components of photosynthesis such as *rbcL*, encoding the large subunit of RuBisCo. These adjustments involve the action of various regulatory proteins, including MRL1, a pentatricopeptide repeat (PPR) protein known to target the *rbcL* 5’-UTR and to control *rbcL* mRNA levels, by preventing RNA degradation (2). Another factor associated with *rbcL* mRNA is the chloroplast ribonucleoprotein (cpRNP) CP29A (3). Proteins of the cpRNP family are characterized by two RNA recognition motifs (RRM) connected by a linker domain. They exhibit high affinity for U/G homopolymers, but lower affinity for cytosine (C) and no affinity for adenine (A) homopolymers (Li and Sugiura, 1991). The interaction of cpRNPs with chloroplast RNAs was examined through a series of in vitro and in vivo tests. Intron-lacking tRNAs and rRNAs were found to have little or no affinity for cpRNPs (4). In contrast, cpRNPs were found to associate with various mRNAs and some intron-containing tRNAs (3–5). A highly enriched transcript in IPs of CP29A was *rbcL* (3). Beyond the identification of co-precipitating transcripts, no details on binding sites in vivo nor whether binding of RNA is direct in vivo are known.

Crosslinking followed by immunoprecipitation (CLIP) employs UV light to create covalent bonds, preserving direct protein-RNA interactions (6–8). Such non-reversible bonds allow harsh washing conditions thereby reducing false positives and facilitating the identification of specific RNA targets. Another advantage of CLIP-based protocols is the higher resolution and thus the ability to identify target sites rather than target RNAs. As the molecular function of RBPs is highly dependent on their binding position within target transcripts, knowledge of binding sites can provide helpful information with respect to an RBP’s biological role. Therefore, a CLIP-based approach was established for the investigation of the molecular role of CP29A in organellar gene expression.

## Results

### eCLIP demonstrates that CP29A binds RNA directly *in vivo* and has a preference for U-rich sequence elements

Only a few in vivo cross-linking and immunoprecipitation experiments, followed by next-generation sequencing (CLIP-Seq) have been conducted in plants (7, 9, 10), and none in chloroplasts. We adapted the eCLIP protocol (11, 12) for chloroplasts, employing specific antibodies for CP29A and CP33B, another cpRNP family member. The binding pattern of CP33B, revealed through prior RIP-Chip results, differs substantially from CP29A (3, 13, 14). Analyzing two RBPs with different target specificities in parallel can unveil potential bias and false-positives in the procedure.

We used 10 billion chloroplasts per CLIP sample, subjecting them to UV crosslinking, followed by lysis and immunoprecipitation with CP29A and CP33B antibodies. Successful precipitation of protein-RNA complexes was confirmed via western blot and radio-labeling of bound RNA (Figure S1A,B). The RNA-protein complexes, larger than the ribonucleoprotein by itself, exhibited size variation visible as a smear on the autoradiogram above the respective cpRNP size (Figure S1B). The corresponding blot section was excised, treated with protease to release RNAs from the nitrocellulose, and the released RNA was processed for library preparation compatible with Illumina sequencing, enabling high-throughput analysis.

Analysis of the CP29A CLIP dataset revealed 69 peaks in 15 chloroplast mRNAs and two tRNAs (Figure 1A). CP33B CLIP identified 52 peaks among 5 mRNAs and one tRNA (Figure S2A). Importantly, no sequence-level overlap was observed between significantly enriched peaks in the CP29A and CP33B datasets, indicating distinct specificities. CP33B’s main target was the *psbA* mRNA, whereas *rbcL* mRNA was a strong target of CP29A (Figure 1A,B, Table S1), consistent with previous analyses (3, 13, 14).

**Figure 1:**
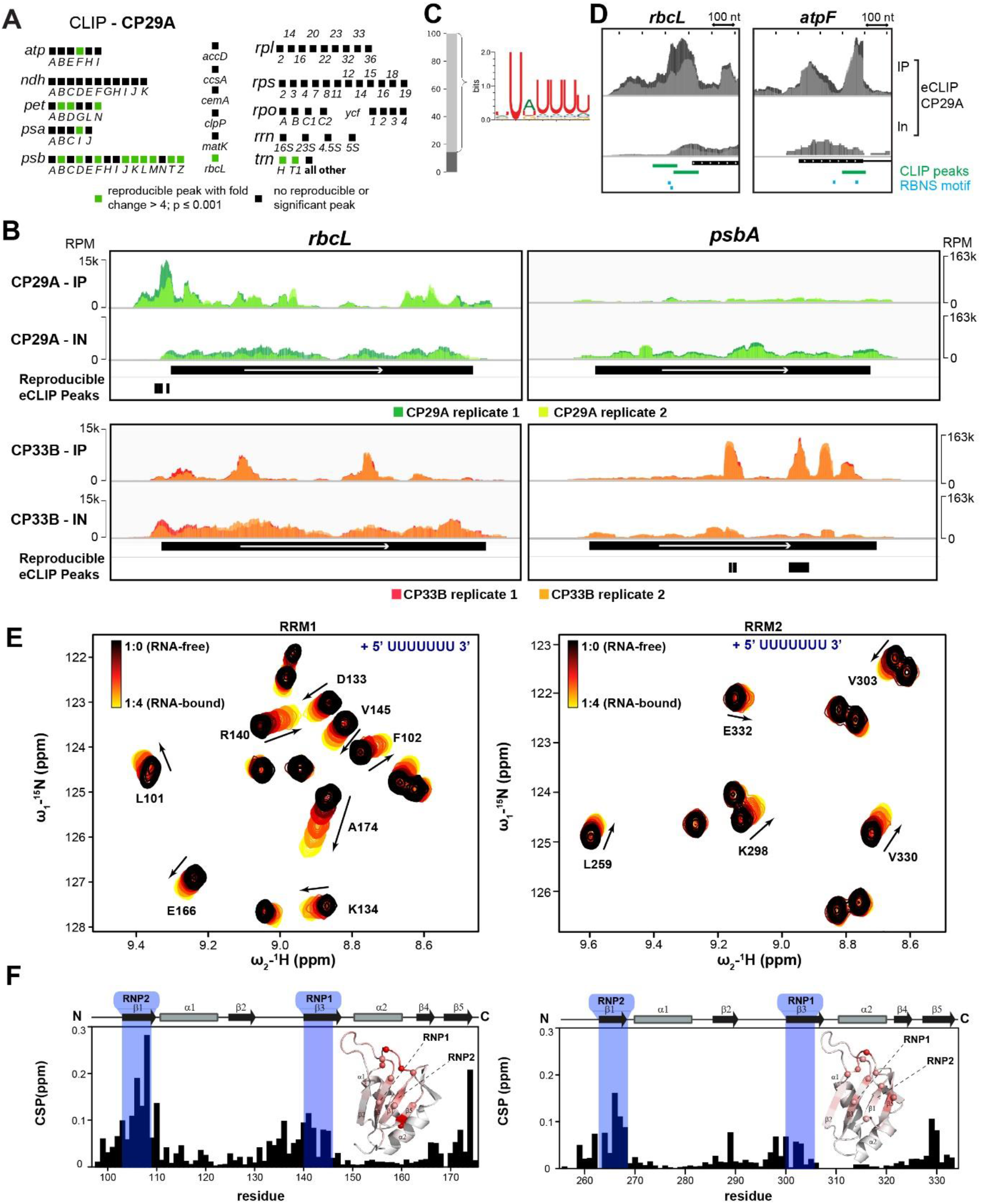
RNA binding analysis of CP29A. eCLIP libraries (IP) and corresponding size-matched input (IN) libraries were prepared from UV-crosslinked chloroplasts. Identified peaks were considered significant and reproducible if fold-enrichment was > 4 and p < 0.001 in eCLIP versus size-matched input, in both replicates. (A) Summary of reproducible eCLIP peaks for CP29A. Detailed CLIP data for CP29A at the (B) *rbcL* and (C) *psbA* locus. Read coverage (in reads per million = RPM) plots are shown for both biological replicates. Reproducible peaks are annotated as black rectangles. (D) Sequence preferences of CP29A were analyzed using the RNA Bind-N-Seq method. Significantly enriched 6-mers were used to construct a sequence logo, which represents most of the observed binding of CP29A. (D) Examples of overlapping RBNS motifs and eCLIP peaks are shown for CP29A in *rbcL* and *atpF*. (E) Overlays of ^1^H-^15^N NMR correlation spectra of the RRM1 (left) and RRM2 (right) domains of CP29A with increasing concentration of “UUUUUUU” RNA oligonucleotide. Chemical shift changes upon RNA binding are highlighted with arrows. (F) Chemical shift perturbations (CSP) for RRM1 (left panel) and RRM2 (right panel) in the presence of 4-fold excess of RNA are shown. The secondary structure of the RRMs is indicated and the RNP motifs highlighted in blue are highlighted. The change in chemical shifts are mapped onto the structural model of individual RRMs with gray to red color gradient.

CP29A peaks were called in all transcriptional units encoding proteins of the photosystem II complex (*psbB-psbT-psbH-petB-petD*; *psbD-psbC-psbZ*; *psbE-psbF-psbL-psbJ*; *psbK-psbI*; *psbM*), except in the monocistronic *psbN* and *psbA* mRNAs (Figure 1A). CP29A also targeted *atpF* mRNA along with monocistronic mRNAs *psaI, petN*, and *rbcL*. Most significant CP29A peaks were within coding sequences, along with peaks in tRNA genes *trnH* and *trnT* and several peaks in 3’ and 5’ untranslated regions (UTRs). UTR regions are known to be of prime importance for the regulation of transcript stability and translation in chloroplasts (15) and are therefore of special interest. Target UTRs of CP29A include the 3’-UTRs of *atpF* (partly overlapping with the CDS), *petN, psbM* and the 5’-UTR of *rbcL* and *petB* (Figure 1B and S3A,B,C).

For CP33B, significant peaks were also predominantly located in the CDS of target genes (Figure 1B, S3E, except for *atpH*, where peaks extended partially into the 3’ UTR (Figure S3E). Besides the top peak in *psbA*, peaks were identified in the second exon of *petB*, in *psbB* and *psbD* mRNA, and in *trnH* (Figure S2A).

It’s essential to acknowledge potential bias in CLIP analysis for CP29A and CP33B, favoring well-expressed genes. All target genes rank in the top two tertiles based on read counts in wild-type Col-0 RNA-seq datasets (16). Consequently, there is a risk of false negatives due to low expression levels, possibly affecting intronic regions. The identified transcripts in CP29A and CP33B datasets through CLIP align with previously identified target mRNAs (3, 13, 14), highlighting the utility of CLIP-based assays in chloroplasts.

### *In vitro* binding data reveal the sequence specificity of CP29A and CP33B

We employed the *in vitro* method RNA Bind-n-Seq (RBNS; 17) to supplement *in vivo* CLIP data. RBNS uses recombinant RBPs to enrich and identify their sequence motifs from a synthesized pool of random RNAs, utilizing high-throughput sequencing. To this end, CP29A and CP33B were genetically modified with an N-terminal glutathione-S-transferase (GST) tag and a streptavidin-binding protein (SBP) tag, expressed in *E. coli*, and purified via the GST tag. After removal of the GST-tag, purified RBPs were incubated at different concentrations (0, 10, 100, and 1000 mM) with a pool of random 40-mer RNA oligonucleotides . After pulldown using the SBP tag, RBP-associated RNAs were isolated. We prepared and sequenced libraries from both the random RNA input pool and the immunoprecipitated (IP) RNAs using Illumina sequencing, resulting in approximately 20 million reads per library. Utilizing the RBNS pipeline (18), we counted all possible k-mers of a specified length in both the IP samples and the input RNA pool and calculated the enrichment of each k-mer as its frequency in the IP samples relative to its frequency in the input pool, denoted as the R-value.

Given that RRM domains typically bind sequences ranging from 2 to 4 nucleotides and that cpRNPs contain two RRM domains, we focused our RBNS library analysis on 6-mers. The highest R-values for both CP29A and CP33B were found in libraries derived from IPs conducted with a protein concentration of 100 nM. Control samples without protein lacked significant 6-mer enrichment. For CP29A, the highest enriched 6-mers were UUUUUU (R=2.26), UUUUUA (R=1.98), and UAUUUU (R=1.97). CP33B’s highest enriched 6-mers were GUUACU (R=3.47), GGUUAC (R=3.39), and GCUACU (R=2.67). These *k*-mers were consistently identified as top *k*-mers across tested protein concentrations. The set of significantly enriched 6-mers of each RBP were used to construct consensus sequence motifs and corresponding position weight matrices (Figure 1C, Figure S2B).

The binding preference of CP29A was most accurately characterized by a degenerate U-rich motif, with A, U, or G being almost equally probable at the third position. The position weight matrix (PWM)-based consensus motif for CP33B corresponded closely to the consensus sequence GHUAUY. We next explored the role of individual RRM domains in binding to U-rich RNA motifs. We monitored the NMR chemical shifts of amides upon titration of a “UUUUUUU” RNA oligonucleotide to the individual ^15^N-labeled CP29A RRM domains. RRM1 exhibited more pronounced chemical shift perturbations (CSPs) upon binding to “UUUUUUU” RNA compared to RRM2 (Figure 1E-F). When mapping the CSP onto structural models of the two RRMs, the two RNP sequence motifs are strongly affected, suggesting that the CP29A RRMs exhibit a canonical interaction with the RNA (Figure 1F). However, the RNA binding affinities estimated from the NMR titrations are different for both RRMs, where RRM2 shows weaker interactions than the RRM1. Overall, RBNS experiments and NMR-based assays of CP29A binding indicate that U-rich sequences are the primary targets of CP29A.To assess the binding of the CLIP derived motifs for CP29a, we performed NMR titrations with short single-stranded DNA oligonucleotides, as we expect that binding specificity of these oligos resembles corresponding RNA ligands (Figure S4A-B). Analysis of the NMR titrations indicated strongest binding of “TGTTTT” and “TTTTTT” oligos to RRM1. On other hand, RRM2 of CP29A shows the strongest response with “TGTTTT” and “GTTACT” oligos (Figure S4C-D). These findings indicate that the RRM1 and RRM2 primarily recognize “T-rich” sequences, mainly “TTT” and “GTT” motifs (Figure S4C), well in-line with our CLIP and RBNS binding analysis.

To assess the occurrence of the RBNS-derived consensus motifs within CLIP peaks versus control peaks, we utilized the Analysis of Motif Enrichment (AME) tool from the MEME suite (19). CLIP-derived peaks within 10 base pairs were merged and extended to match the largest peak size (53 nucleotides for CP29A, 82 for CP33B). The control peak set was created by shuffling all chloroplast sequences used in the CLIP analysis, ten times, resulting in ten random versions for each gene. From these, 1000 peaks, matching the largest CLIP peak size, were used as a control set for the analysis of each RBPs motif. Additionally, chloroplast gene sequences were employed to establish a 0-order background model for the AME tool.

The CP29A PWM-based RBNS motif was statistically significant (adjusted p-value < 0.05) enriched in 13 of 23 CLIP-derived merged peaks. for example in *atpF* (Figure 1D). Notably, this motif appeared twice in the *rbcL* 5’-UTR CLIP peak, which exhibited the lowest p-value in the peak enrichment analysis (Figure 1D). In contrast, CP33B AME analysis did not reveal significant enrichment of the RBNS motif in merged CLIP peaks. However, this motif was found in 5 of the 8 merged peaks, including the *petB* and the *psbA* CLIP peak, which had the lowest p-value in the CP33B CLIP analysis (Figure S2C).

The correlation between the sequence preferences of CP29A and CP33B identified by RBNS and their respective *in vivo* binding sites revealed through CLIP provides a foundation for establishing a high-confidence set of binding sites for both proteins.

### CP29A binds adjacent to the predicted binding sites of the PPR protein MRL1 in the *rbcL* 5’-UTR

The primary binding site for CP29A was identified in the *rbcL* 5’UTR (Figure 1B, Table S1). A crucial factor known to influence the expression of *rbcL* is the pentatricopeptide repeat (PPR) protein MRL1 (2). PPR proteins are known to produce short RNA fragments that are protected from nuclease degradation due to their tight binding (20–22). We previously discovered a short RNA fragment, designated as C28, which precisely matches the end of the mature, 5′-processed rbcL mRNA (20, 21). The formation of the processed 5’ end of the *rbcL* RNA is dependent on the PPR protein MRL1 (2).

To assess MRL1’s role in *rbcL* 5’ sRNA/C28 sRNA accumulation, we performed small RNA sequencing on the Arabidopsis *mrl1* mutant, comparing it with wild-type (wt) plants. Our analysis identified C28 as the sole small RNA (sRNA) exhibiting differential accumulation across the plastid transcriptome (Figure 2A,B). Validation via RNA gel blot hybridization supported this finding (Figure 2C). Control mutants lacking the PPR protein HCF152 or the H-TRP protein HCF107 accumulated the *rbcL* 5’ footprint with a slight signal reduction (Figure 2C). This decrease was attributed to pleiotropic effects stemming from their photosynthetic defects. These controls substantiate the specificity of C28 loss observed in *mrl1* mutants.

**Figure 2:**
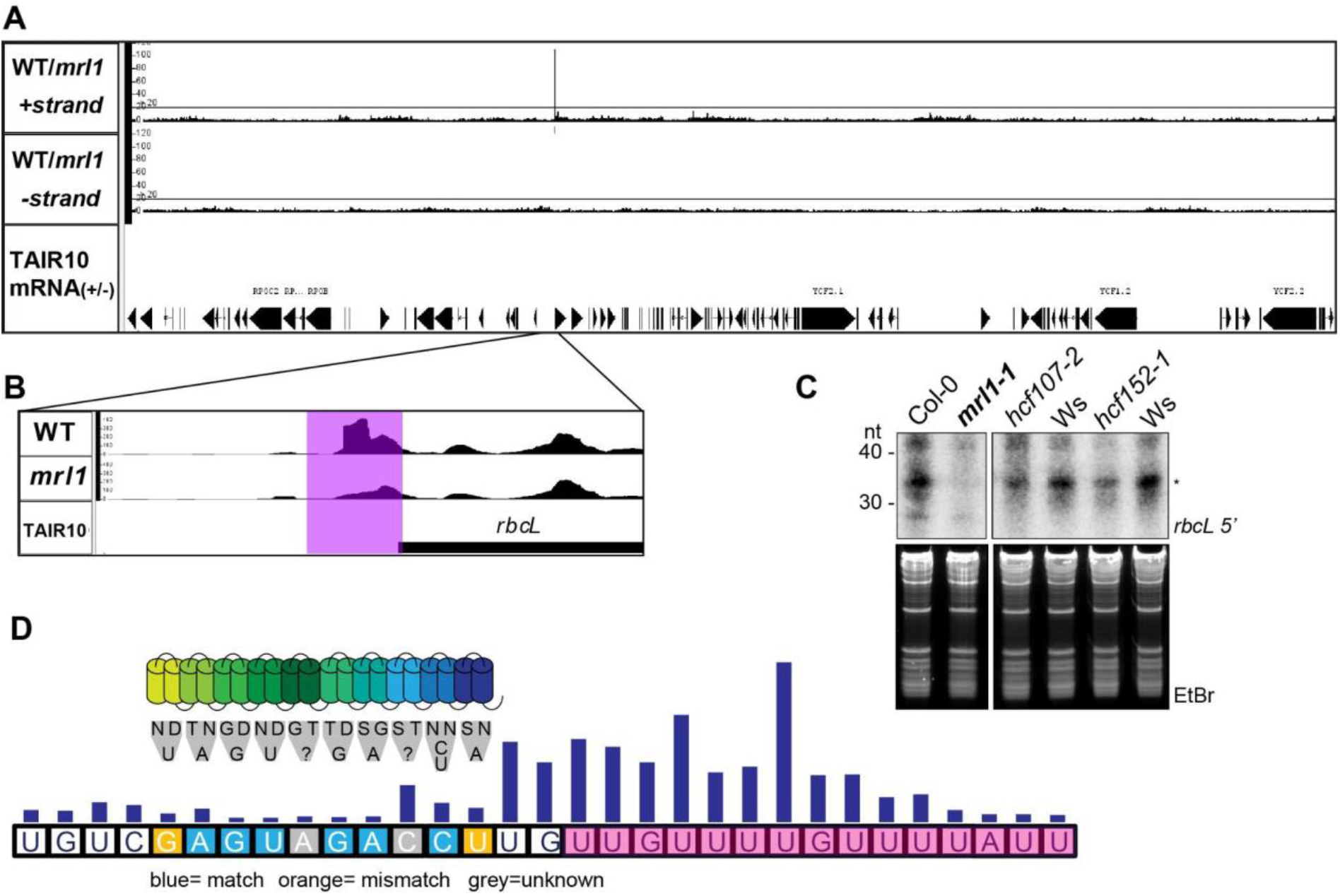
Identification of differential small RNA accumulation in *mrl1* mutants. (A) Ratios of wt and mutant small RNA sequencing counts. Ratios for the positive and negative strand are shown for the 5’ end of small RNAs. Only for the sRNA C28 in the *rbcL* 5’UTR, a high ratio of reads in wt versus *mrl1* was identified. (B) An sRNA coverage graph of the corresponding genomic region is shown here. The area of differential RNA accumulation is shaded in violet. (C) Small RNA gel blot analysis of C28 in *mrl1* mutants and the corresponding wt were grown on soil for three weeks. The *high chlorophyll fluorescence* (*hcf*) mutants *hcf107-2* and *hcf152-1* were analyzed as controls with known sRNA accumulation defects. Since these mutants are in the Wassilewskija (Ws) background, Ws plants were used as control. (D) Mapping of the predicted MRL binding site, CP29A eCLIP data and CP29A RBNS data onto the *rbcL* sRNA C28 sequence. The sequence of C28 is shown with framed capital letters. MRL1 has 10 PPRs, which are schematically shown as an array of two joined helices. The key amino acids are shown below together with their predicted base target. CP29A eCLIP-coverage data of read termini are shown as a bar graph above the bases in the sRNA sequence. The sequences matching the RBNS consensus shown in Figure 2 are highlighted in magenta.

The binding site of MRL1 within the *rbcL* mRNA was hypothesized to be near its 5’-end (2). Using the PPR code (23), we predicted MRL1’s binding sequence based on the specific amino acids in each repeat of its PPR tract to align partially with the 5’-*rbcL* sRNA (Figure 2D), corroborating the region proposed earlier (2). However, the sRNA extends considerably beyond this binding site. Interestingly, when analyzing the CP29A CLIP-data, we noticed that one peak is close to the MRL1-dependent C28 sRNA. Reads from eCLIP terminate at the protein-RNA cross-link site, providing base-specific resolution. Mapping read ends showed a peak downstream of the predicted MRL1 site, still within the C28 sRNA (Figure 2D). This alignment is notable as the U-rich target sequence in the *rbcL* peak aligns with the consensus sequence from our RBNS analysis (Figure 1C, Figure 2D). This suggests a potential functional overlap in the binding regions of MRL1 and CP29A within the *rbcL* mRNA, with CP29A binding downstream of the MRL1 site.

### Arabidopsis CP29A mutants do not show a change in *rbcL* mRNA levels nor is translation of the *rbcL* mRNA reduced

Considering MRL1’s role in stabilizing and translating *rbcL* (2), we studied *rbcL* expression in *cp29a* null mutants under standard and cold acclimation conditions. We treated the plants for three days with 12°C, conditions that do not yet lead to a photosynthetic or macroscopic phenotype, thus avoiding secondary effects due to failed chloroplast biogenesis (16). We found no significant alterations in *rbcL* mRNA accumulation or RbcL protein levels between wild-type and *cp29a* mutants under either condition (Figure S5A-D). As RuBisCo has a long half-life (7 days; 24, 25) smaller changes on the level of translation activity may not become visible after three days on the backdrop of the large pool of stable RuBisCo protein. We therefore next tested the synthesis of chloroplast proteins by labeling with 35S methionine. After three days of cold exposure, no significant alterations were detected in protein synthesis of either RbcL or D1 as a control protein in the chloroplasts of the *cp29a* null mutants (Figure S5E-H). These findings suggest that CP29A absence does not notably affect *rbcL* mRNA accumulation or translation in short-term cold-treated *Arabidopsis* tissue.

### CRISPR-Cas9-induced null mutants of tobacco CP29A show photosynthetic deficiency in the cold

*Arabidopsis thaliana*, a cold-adapted species, thrives in temperate zones and can grow even at 4°C. We asked whether the disruption of a CP29A relative in tobacco (*Nt*CP29A), a tropical plant, which does not often need to acclimate to cold, might yield similar outcomes. This choice was informed by evidence showing that *Nt*CP29A localizes to the chloroplast, can bind *rbcL* and other RNAs *in vitro* and *in vivo* (4, 26), and stabilize RNAs in vitro (27), mirroring some of the functions of *CP29A* in *A. thaliana*.

Tobacco, an allotetraploid derived from *N. sylvestris* and *N. tomentosiformis*, harbors two sets of alleles denoted here as *NtsCP29A* and *NttCP29A*. Utilizing CRISPR/Cas9 mutagenesis, we targeted all four alleles to generate null mutants (Figure S6A). In the T1 generation, we obtained plants lacking either *NtsCP29A* or *NttCP29A*, but not plants with large deletions in both alleles. After self-fertilization and crossing, we generated a homozygous *Ntscp29a*/*Nttcp29a* line, confirmed by PCR (Figure S6B) and sequencing (Figure S6C). We designate these plant lines as *Ntcp29a* mutants.

Under standard conditions, Ntcp29a plants resembled wild type, showing no abnormalities (Figure S6D). After 14 days of low temperatures, they still displayed no macroscopic defects (Figure S6D). Given that *Atcp29a* mutants experience photosynthetic impairments under prolonged cold exposure (16), we evaluated the quantum yield efficiency of photosystem II in *Ntcp29a* mutants. The Fv/Fm values were slightly decreased under normal conditions, worsening significantly in the cold compared to wild type (Figure 3A,B). This decline resembles observations in *Arabidopsis* mutants. Notably, while *Atcp29a* mutants showed reduced Fv/Fm only in young leaf tissue (Figure 3C), *Ntcp29a* mutants exhibited decreased efficiency across entire leaves under both conditions (Figure 3A)

**Figure 3:**
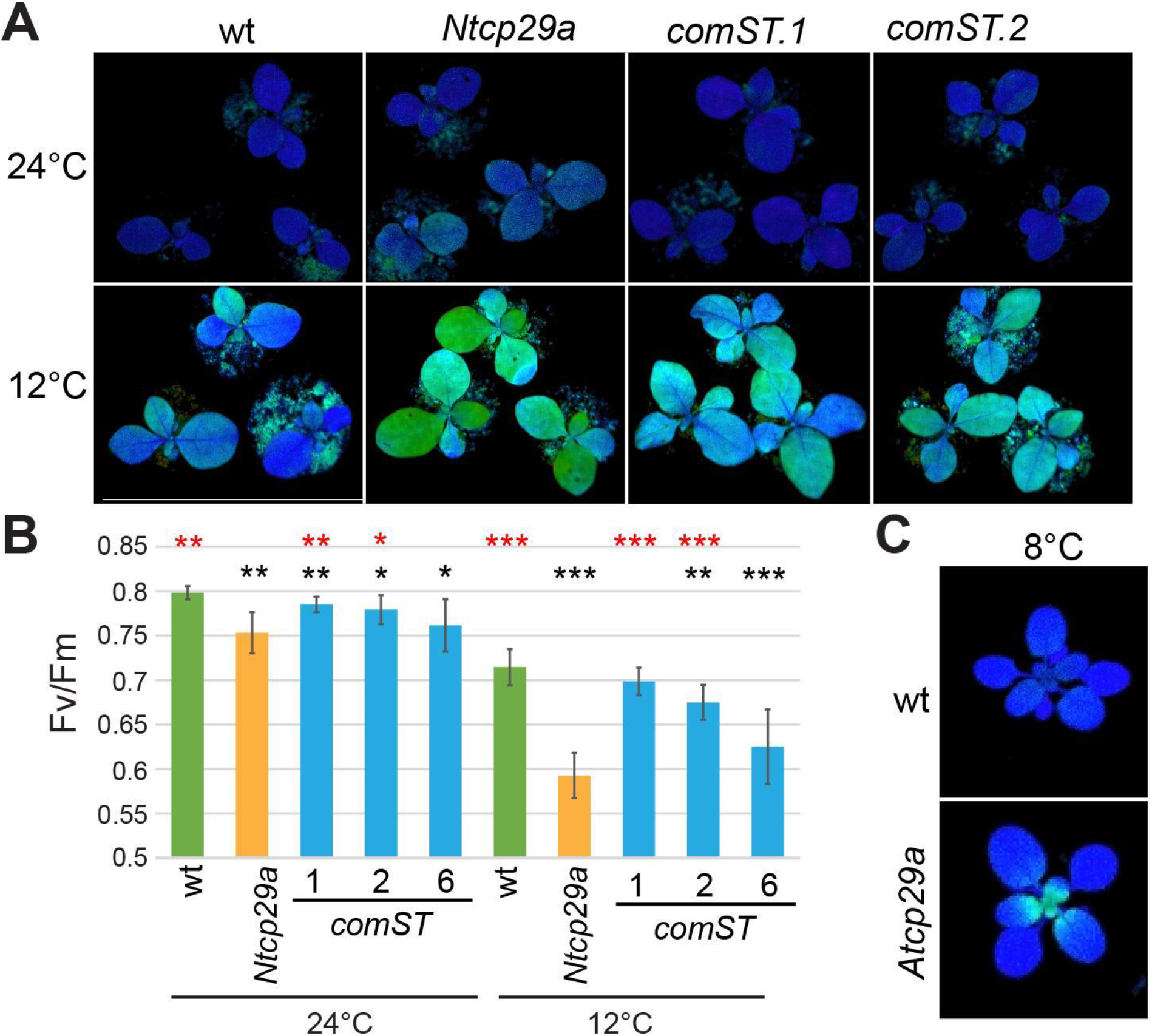
Chlorophyll a fluorescence analysis of *Ntcp29a* mutants. (A) Visualization of the maximum quantum yield of photosystem II (FV/FM) with an imaging PAM under normal growth conditions and after cold treatment. Wt, *Ntcp29a* mutants and three independent complementation lines (*comST*) were grown for 21 days at 24°C and then transferred to 12°C for seven days. (B) Fv/Fm values based on the measurements of eight individual plants for each plant line. Bars indicate the standard deviation and the asterisks represent the statistical significance (*p-value < 0,05; **p-value < 0,01; ***p-value < 0,001), as evaluated by Student’s t-test. red asterisks = comparison to *Ntcp29a* mutants; black asterisks = comparison to wt. Phenotype of Arabidopsis *Atcp29a* mutants in comparison to wt after 14 days at 21°C and 10 days at 8°C.

We confirmed that the absence of *NtCP29A* caused the photosynthetic defect by a complementation test on the null mutant. Introduction of full-length genomic versions of both *N. sylvestris* and *N. tomentosiformis* alleles via *Agrobacterium*-mediated transformation resulted in restored photosynthetic activity (Figure 6A,B). Recovery levels varied across complementation lines, likely due to varying transgene expression levels. Overall, these analyses confirm that the tobacco *NtCP29A* is essential for maintaining full photosynthetic performance during cold acclimation.

### Tobacco CP29A mutants have reduced levels of *rbcL* mRNA and RbcL protein in the cold

To identify the molecular cause of the diminished photosynthetic performance in *Ntcp29a* mutants, we examined chloroplast RNA processing and accumulation under normal growth conditions (21 days at 24°C) and after exposure to cold (21 days at 24°C +7 days at 12°C). RNA extraction was done from one half of the fourth primary leaf, followed by Illumina-based RNA-seq after rRNA depletion. Protein analysis was done on the other half of the leaf (details below).

Considering that *Atcp29a* mutants exhibited splicing defects after short-term cold exposure before visible bleaching (16), we investigated RNA splicing in the tobacco mutants. Using the Chloro-Seq pipeline (28), we assessed the splicing efficiency of the 21 introns in the tobacco chloroplast genome. Our results showed no significant differences in splicing efficiency between the two genotypes under either normal or cold conditions (Figure S7A,B). In addition, we quantified chloroplast RNA editing in wt and mutants under both conditions, again showing no significant changes (Figure S7C,D).

Next, we investigated RNA accumulation differences between the two genotypes under normal and cold temperatures. Sequencing results were analyzed using the DeSeq2 pipeline (29). Principal component analysis (PCA) showed good clustering of replicates (Figure 4A), with the first principal component differentiating growth conditions, while the second, much smaller component reflected the variance between mutant and wt samples (Figure 4A). This aligns with the expected impact of cold on RNA pools, resulting in marked differences in transcript accumulation.

**Figure 4:**
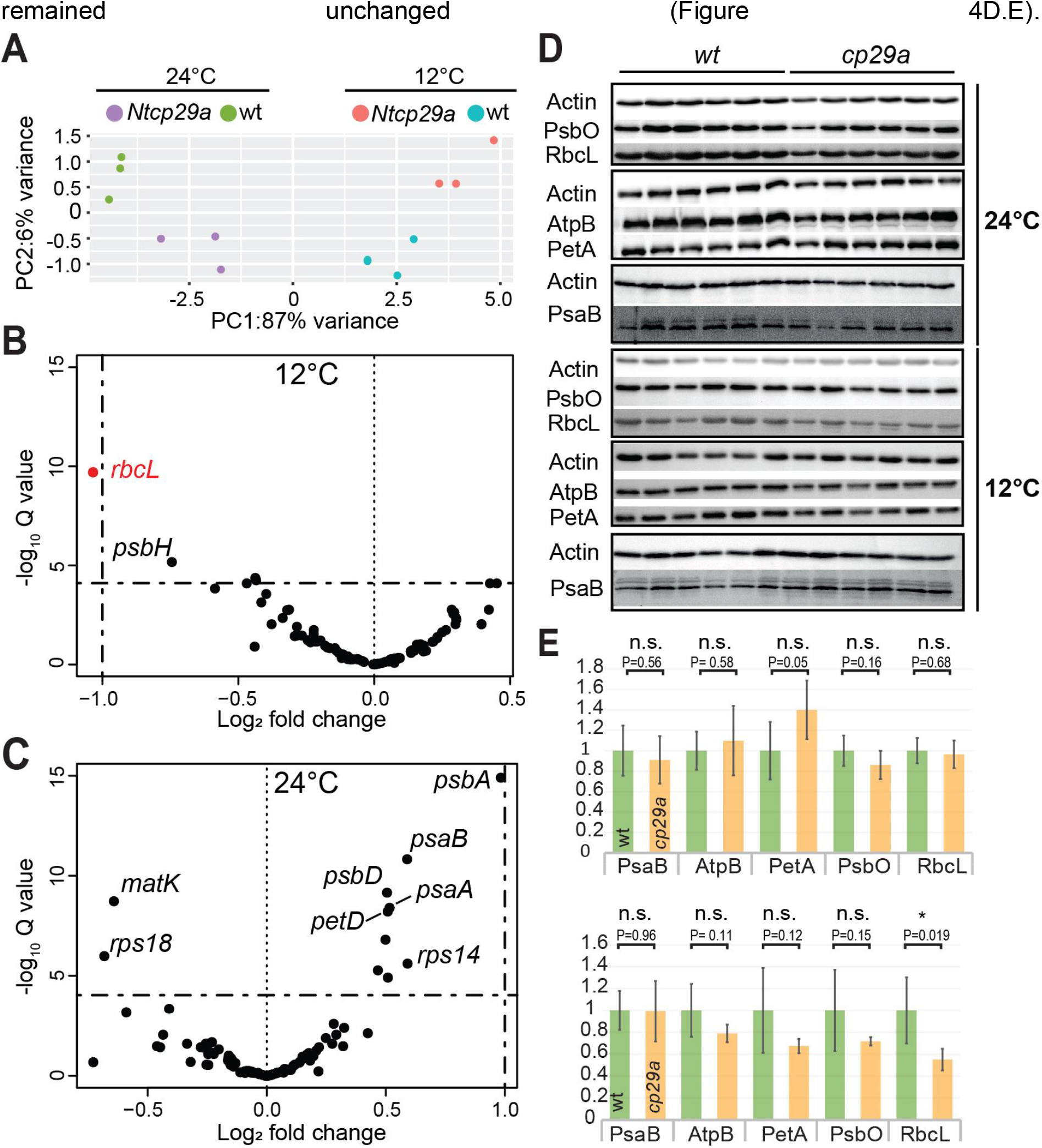
Analysis of *rbcL* expression in *Ntcp29a* mutants. (A) Principal component analysis of *Ntcp29a* mutant and wt leaf RNA-Seq samples (n=3). Plants were grown for 21 days at 24°C and then directly analyzed or transferred for 7 days to 12°C and then analyzed. (B) Volcano plot of RNA-Seq results for chloroplast RNAs from cold-treated mutant versus wt plants after cold treatment. Only the *rbcL* mRNA shows a more than twofold downregulation in the cold, which is statistically significant (p<0.001; significance test: wald test adjusted with Benjamini Hochberg). Dashed lines show selected thresholds for fold change and Q value. (C) As in (B), but for plants grown at standard growth temperature, 24°C. No expression changes above the twofold threshold were detected. (D) Immunoblot analysis of leaf extracts of tobacco seedlings. Samples from plants analyzed for RNA accumulation in A-C were subject to immunological analyses. A total of six wt and six *Ntcp29a* mutant plants were analyzed. (E) Quantification of immunoblot analysis from (D). * = p<0.05; student’s t-test.

We compared chloroplast gene expression in *Ntcp29a* mutants to wild type (wt), focusing on normalized RNA coverage fold changes. Out of 94 genes meeting detection criteria, *rbcL* showed the strongest decrease, reaching 49% of wt levels, exclusively under cold conditions (Figure 4B,C). Expression changes occurred also in other genes, including eCLIP targets of CP29A, but none reached our fold-change threshold. Overall, *Ntcp29a* mutants exhibited a significant and specific reduction in *rbcL* mRNA at low temperatures.

We next investigated if the decrease in *rbcL* mRNA affected RbcL protein accumulation by immunoblot analyses. Apart from RbcL, we examined other subunits from the Photosystem I and II, the ATPase, and the cytochrome b6f complex. Only RbcL showed a significant reduction specifically in the cold in *Ntcp29a* mutants, to less than 50% of wt levels, while other proteins

## Discussion

### eCLIP reveals a post-transcriptional operon rich in PSII mRNAs for AtCP29A

In this study, we employed for the first time the eCLIP-Seq method to investigate RNA interactions of a chloroplast RNA-binding protein (RBP). We demonstrate that despite high pigment concentrations in chloroplasts UV-cross-linking is efficient enough to allow precipitation of RBP:RNA complexes. Our analysis revealed highly specific, distinct peaks, distinguishing between closely related cpRNPs, thereby enabling specific ligand identification for *AtCP29A*. This approach offers significant advantages over previous methods like RIP, which might capture larger RNA-protein complexes, obscuring specific interactions (3, 4).The eCLIP-based analysis corroborated previous findings, identifying CP29A as a chloroplast RNA-binding protein (RBP) with a multitude of target transcripts. This includes the reconfirmation of previously identified targets like *rbcL, psbD*, and *psbB*. Other previously identified targets were not identified in the current eCLIP analysis. This discrepancy could be attributed to the absence of fixed cut-off criteria in the earlier RIP-Chip experiments, suggesting that these genes might represent minor targets of CP29A. They possibly did not meet the more stringent threshold criteria applied in the present eCLIP experiment.

Many of these binding sites are in mRNAs coding for photosystem II subunits as well as for other photosynthetic electron transport components. For example, within the *psbB* operon, genes like *psbB, psbT, petB*, and *petD* were identified as targets. Similarly, in the *psbE* operon, *psbF, psbL*, and *psbJ* emerged as targets. This finding is particularly noteworthy because the input RNA in the eCLIP-based approach undergoes fragmentation during cell lysis, and the resulting library is size-selected. This process allows for the differentiation of binding sites even within the same gene, as exemplified by the identification of distinct binding sites in the 5’ UTR and exon 2 of *petB* (Figure S3B). Additionally to polycistronic photosystem II transcripts, the functionally related monocistronic transcripts of *psbM, psbK*, and *petN* were also identified as targets. The preferential association of AtCP29A with these functionally related transcripts suggests the potential existence of a post-transcriptional operon, a concept previously described in various species (31). Another cpRNP, AtCP31A has been demonstrated to associate preferentially with mRNAs encoding subunits of the NDH complex (32), which are not targets of AtCP29A. AtCP33B with its preference for the *psbA* mRNA that is neither targeted by AtCP29A nor by AtCP31A has yet another non-overlapping target range. Therefore, the data presented here suggest that cpRNPs might serve as operon-defining chloroplast RNA-binding proteins, indicating a sophisticated level of post-transcriptional regulation within the chloroplast.

While our study identified an intriguing target profile for AtCP29A, most of these mRNAs did not exhibit significant accumulation changes in previous RNA-Seq analyses conducted on *Atcp29a* mutants (16), nor in the RNA-Seq analysis of *Ntcp29a* mutants. However, many AtCP29A binding sites are within coding regions, suggesting a role in translation. Supporting this, Ribo-Seq analysis indicated reduced translation in *Atcp29a* mutants under cold conditions (16). Thus, while CP29A’s direct impact on mRNA accumulation seems limited, its potential role in translation regulation or other subtle RNA processing aspects in *cp29a* mutants merits further investigation, especially concerning plant responses to cold stress. Recently, we demonstrated that CP29A can form granular structures at low temperatures through phase separation, which affects the splicing of specific chloroplast introns (16). Whether granule formation also plays a role for *rbcL* accumulation in tobacco remains to be determined.

### Sequence Preferences of AtCP29A

The RBNS method unveiled the sequence preferences of cpRNPs, challenging previous notions of their nonspecificity (33). It unveiled consistent k-mer enrichments across all libraries associated with a specific cpRNP but not in the zero-protein control, validating RBNS specificity using recombinant cpRNP proteins. The identification of distinct target sequence motifs for CP29A and CP33B indicates that, although cpRNPs bind to many chloroplast RNAs, their binding is specific and not merely due to a general affinity for RNA. Notably, CP29A’s enrichment of polyU motifs aligns with previous findings on tobacco homologs’ U-binding ability (34). Since both RRM domains show a preference for U-rich sequences in our NMR-based RNA binding analysis, we speculate that CP29A can bind either longer U-stretches on single transcripts or can bridge two separate RNAs with U-rich motifs. Binding of non-consecutive RNA motifs has been indeed observed in RBPs with multiple RRM domains (35, 36). The latter could be important for network formation during the described phase separation of CP29A, which depends on the prion-like linker domain between CP29A’s two RRMs, but likely also requires the two RRM motifs themselves (16).

Extensive profiling of human RNA-binding proteins (RBPs) revealed a preference for low-complexity target motifs (18). Given that chloroplast genomes typically have low GC-content, the predominance of simple U or A binding motifs in chloroplast RBPs might represent an evolutionary adaptation. Such a motif could be functionally significant to co-regulate multiple transcripts post-transcriptionally. In contrast to CP29A, CP33B binds to a more complex GHUAUY motif, potentially explaining its fewer target RNAs compared to CP29A and CP31A. This suggests that the complexity of binding motifs may influence the range of target RNAs an RBP can regulate.

### NtCP29A supports *rbcL* mRNA accumulation in the cold in tobacco

RNAs often interact with multiple RBPs that work together to regulate the expression or repression of their target transcripts. Many RBPs with RRM domains, such as twin RRM-proteins in the heterogeneous nuclear RNP (hnRNP) class, function in coordination with other RBPs (37). In this context, it was observed that a high-confidence eCLIP binding site on the *rbcL* mRNA in chloroplasts is proximal to a region proposed for MRL1 protein targeting. Genetic analyses of *mrl1* mutants in *Chlamydomonas reinhardtii* and *Arabidopsis* (2), alongside predictions of MRL1-binding sites and identification of MRL1-dependent *rbcL* sRNA accumulation, imply MRL1 binding to the *rbcL* mRNA 5’-end near the AtCP29A binding site. In tobacco, *rbcL* mRNA levels were decreased by 50% in the cold, paralleling RbcL protein reduction and implicating CP29A as a critical RuBisCo expression factor in cold conditions. Given CP29A’s location downstream of the MRL1 binding site, it’s hypothesized that CP29A might assist MRL1 in stabilizing the *rbcL* mRNA against exonucleolytic degradation (2), possibly by aiding MRL1 to bind its target sequence. PPR proteins like MRL1 favor single-stranded RNA targets, and 5’-UTR secondary structures are known to hinder chloroplast mRNA translation (38, 39). Cold stabilizes RNA secondary structures, raising speculation that CP29A might facilitate MRL1-*rbcL* 5’-UTR association in the presence of potentially detrimental RNA structures. Analysis of the *rbcL* mRNA structure under varied temperatures and in *cp29a* mutants could further test this hypothesis.

The anticipated outcome of a 50% reduction in RbcL, and consequently in RuBisCo, is a decrease in carbon fixation. Mutants with alterations in the catalytic center of RuBisCo (center is encoded by *rbcL*), as well as *rbcL* deletion mutants, exhibit pronounced photosynthetic deficiencies and associated growth defects (40, 41). Hypomorphic RuBisCo mutants caused by lowered RbcS levels show reduced Fv/Fm values similar to Ntcp29a plants, even at normal temperatures (42). Comparatively, plants grown in low temperatures typically exhibit higher concentrations and activities of RuBisCo and other enzymes involved in photosynthetic carbon metabolism than those grown at higher temperatures (43–46). This is supported in *Arabidopsis* by an increase of *rbcL* mRNA in the cold and a switch of nuclear-encoded RbcS gene isoforms (42, 47). Since *rbcL* is the rate-limiting factor for RuBisCo assembly (48), *rbcL* expression needs to be sustained or possibly increased in cold environments. The basis for cold-driven expression of RuBisCo was so far unclear, but our data suggest that CP29A plays a role here, at least in tobacco.

## Materials and Methods

The Supplementary Information (SI) Appendix provides comprehensive details on the Materials and Methods utilized in this study. Within the SI.I, a thorough protocol for the chloroplast eCLIP method is outlined, along with specifics of the computational analysis performed. Details pertaining to RBNS are also included in SI.I. Information regarding the NMR analysis can be found in Section II of the Supplementary Information. Section III presents the CRISPR/Cas9 workflow that was followed. Furthermore, Section IV details the growth conditions and standard techniques employed in this study.

## Data Availability

All study data are included in the article and SI Appendix. RBNS and eCLIP next generation sequencing data associated with this manuscript can be found at SRA, BioProject PRJNA1043672. Reviewer access: https://dataview.ncbi.nlm.nih.gov/object/PRJNA1043672?reviewer=ik5egv4bl29p1mun39ojfin73v

RNA-Seq data have been deposited in the National Center for Biotechnology Information Gene Expression Omnibus (GEO) repository (GSE248582). To review GEO accession GSE248582: Go to https://www.ncbi.nlm.nih.gov/geo/query/acc.cgi?acc=GSE248582. Enter token uzavwaowhzanvyv into the box.

## Acknowledgements

The vectors for CRISPR/Cas9 mutagenesis were kindly provided by Kerstin Kaufmann (HU Berlin). We are grateful to C. Stock (HU Berlin) for assistance in cultivation of tobacco tissue, genotyping and immunoblotting. We thank Sam Asami and Gerd Gemmecker for support with NMR experiments. We acknowledge NMR facilities at the Bavarian NMR Center (BNMRZ) at the Technical University of Munich, Germany. We acknowledge SBGrid and the NMRbox server for providing access to NMR softwares. This work was funded by DFG TR175 projects A02 to CSL and A07 to HR. MS and CSL wish to acknowledge funding by DFG SPP1935.

