## Supplemental Figures and Tables for "The Chloroplast RNA Binding Protein CP29A supports *rbcL* expression during cold acclimation"

### Supplementary Information (SI) Appendix

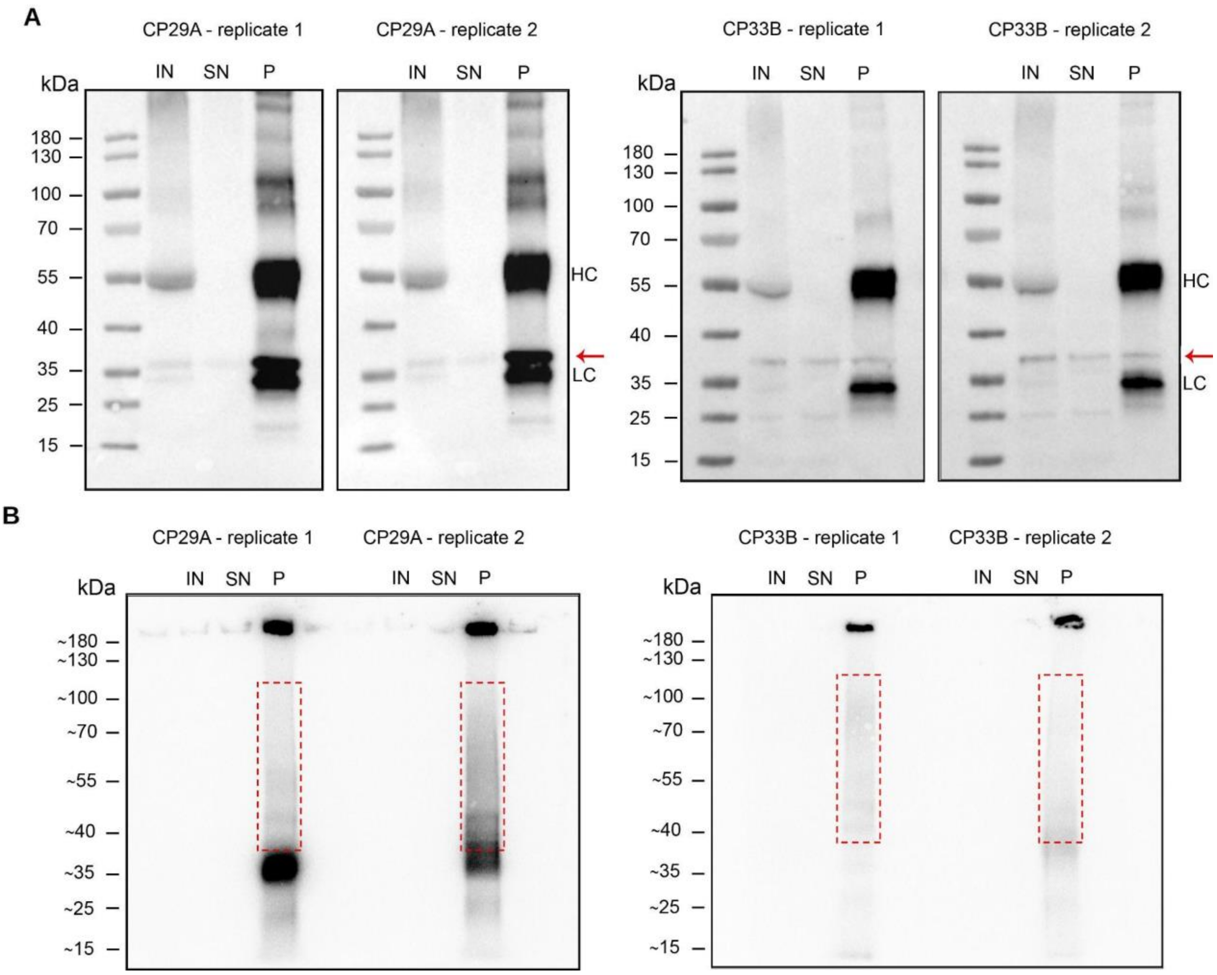


**Figure S1: Monitoring of the immunoprecipitation and crosslinking of CP29A and CP33B during CLIP.**

Samples were taken during the CLIP protocol to verify successful immunoprecipitation and RNA recovery. (A) Samples of the crosslinked chloroplast lysate (IN=input), the supernatant after immunoprecipitation (SN) and the washed beads (P=pellet) were size separated by Bis-Tris PAGE (4-12%). After transfer to nitrocellulose membranes, the blots were probed using the CP29A- and CP33B-specific antibodies, respectively. IgG-HRP was used as secondary antibody. Red arrows indicate the respective cpRNP signal. (HC) IgG heavy chain (LC) IgG light chain. (B) RNAs in the pellet samples were 5’-labeled with [gamma$^{32}$P]-ATP before gel electrophoresis. Radioactive signal was detected by a phosphorimager before the blots were probed with antibodies. The red dashed boxes indicate the size range that was cut from the actual preparative blots (approximately 75 kDa above the respective RBP).

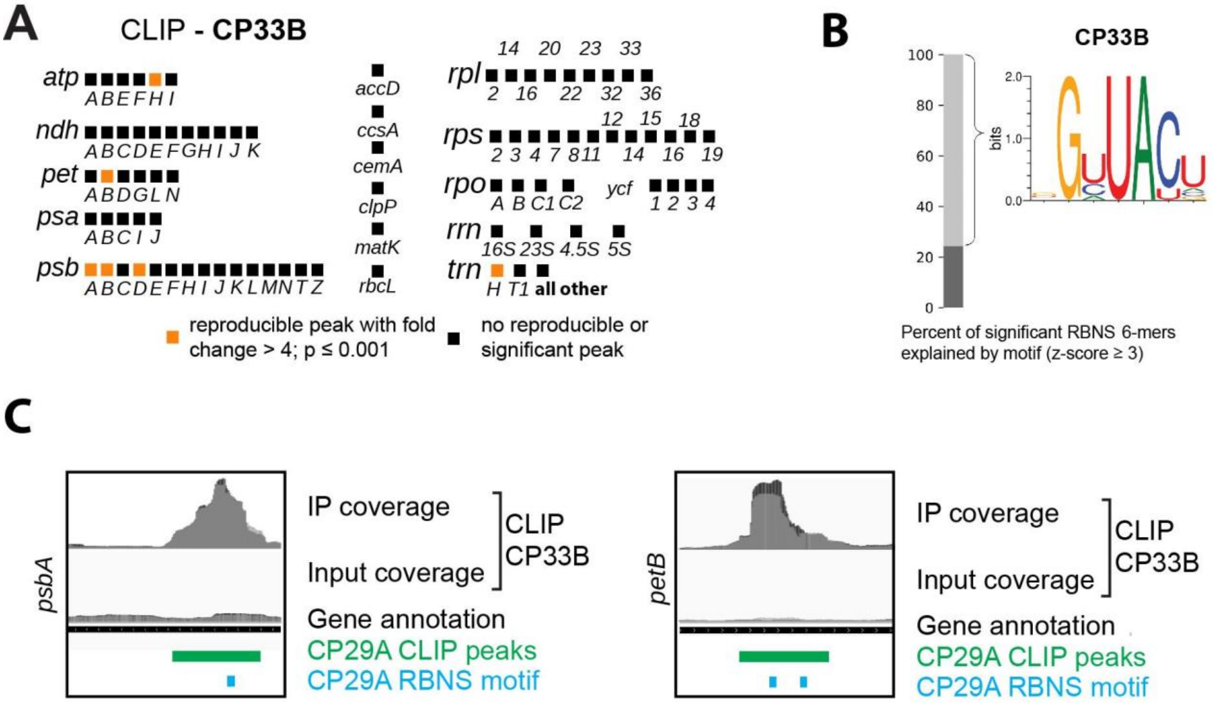


**Figure S2: Binding analysis of CP33B using an eCLIP-derived approach.** CLIP libraries (P) and corresponding size-matched input (IN) libraries were prepared from UV-crosslinked chloroplasts. The library preparation workflow was adapted to chloroplasts based on the eCLIP protocol (12). Data analysis was performed using the eCLIP pipeline (v0.3.99) and the chloroplast genome (TAIR10; customized AtRTD2 annotation with designated 100 nt of 5'- and 3'-UTR). Identified peaks were considered significant and reproducible if fold-enrichment was > 4 and p < 0.001 in CLIP versus size-matched input, in both replicates. (A) Identified reproducible peaks for CP33B. In detail CLIP data for CP33B at the (B) *rbcL* and (C) *psbA* locus. Read coverage (in reads per million = RPM) plots are shown for both biological replicates. Identified reproducible peaks are annotated as black rectangles. (D) Sequence preferences of CP33B were analyzed using the RNA Bind-N-Seq protocol (RBNS; 18). The *in vitro* approach relies on the enrichment of specific sequence elements from a random RNA input pool by purified, recombinant RNA-binding proteins. Significantly enriched 6-mers in RBNS data sets were used to construct a sequence logo, which represents most of the observed binding of CP33B. (E) Examples of overlapping RBNS motifs and CLIP peaks are shown for CP33B in *psbA* and *petB* (D). Coverage graphs (RPM) of normalized CLIP and size-matched input libraries are shown in addition.

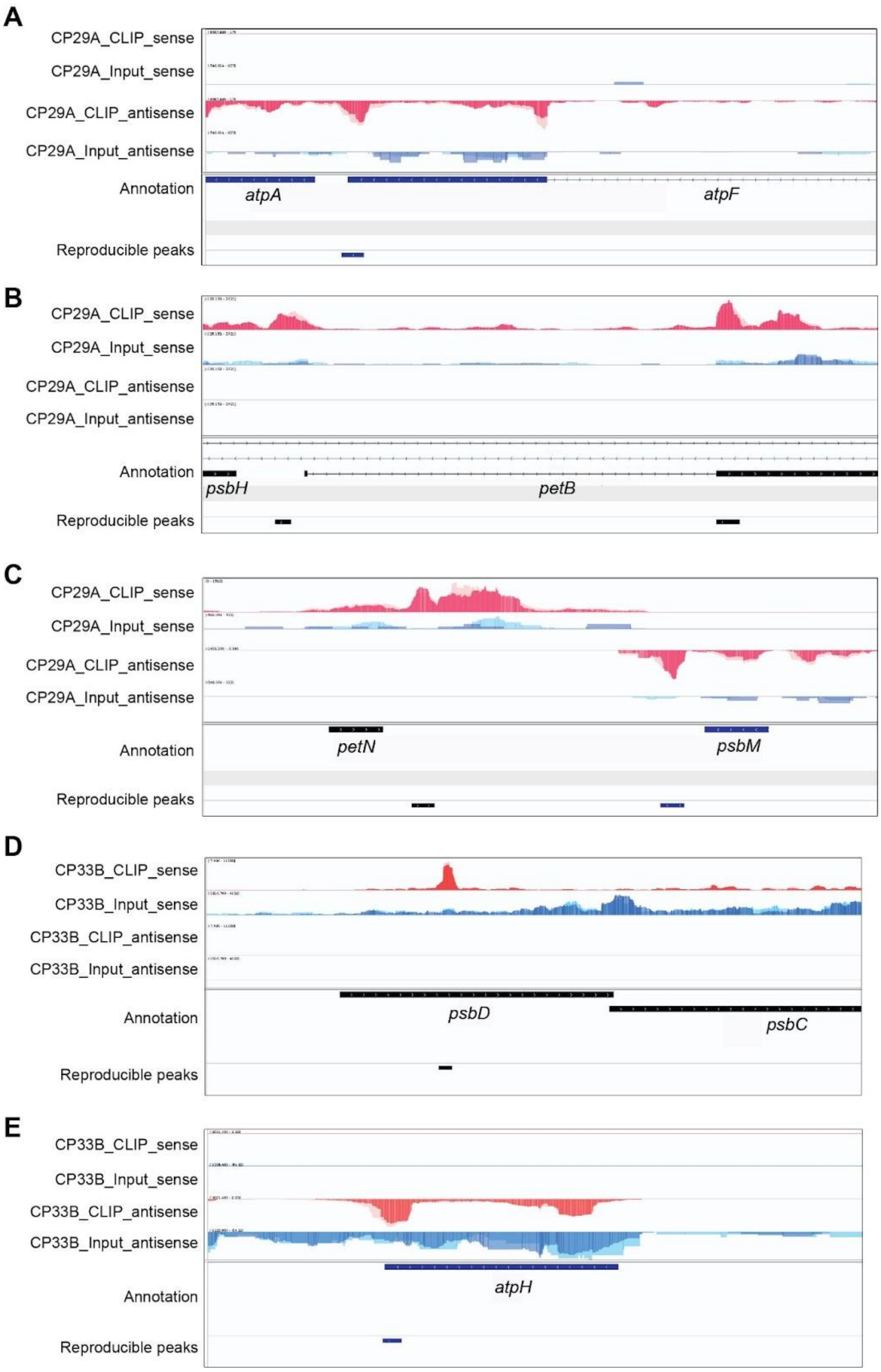


**Figure S3: Binding analysis of CP29A using an eCLIP-derived approach.** CLIP libraries and corresponding size-matched input (smInput) libraries were prepared from UV-crosslinked chloroplasts. The library preparation workflow was adapted to chloroplasts based on the eCLIP protocol (12). Data analysis was performed using the eCLIP pipeline (v0.3.99) and the chloroplast genome (TAIR10; customized AtRTD2 annotation with designated 100 nt of 5'- and 3'-UTR). Identified peaks were considered significant and reproducible if fold enrichment was > 4 and $p < 0.001$ in CLIP versus smInput, in both replicates. (A) Identified reproducible peaks for *atpF*. (B) Identified reproducible peaks for *petB*. (C) Identified reproducible peaks for *petN* & *psbM*. (D) Identified reproducible peaks for *psbD*. (B) Identified reproducible peaks for *atpH*. Coverage graphs (RPM) are shown for both biological replicates (light & dark colors). Identified reproducible peaks are annotated as rectangles.

### Figure S4A

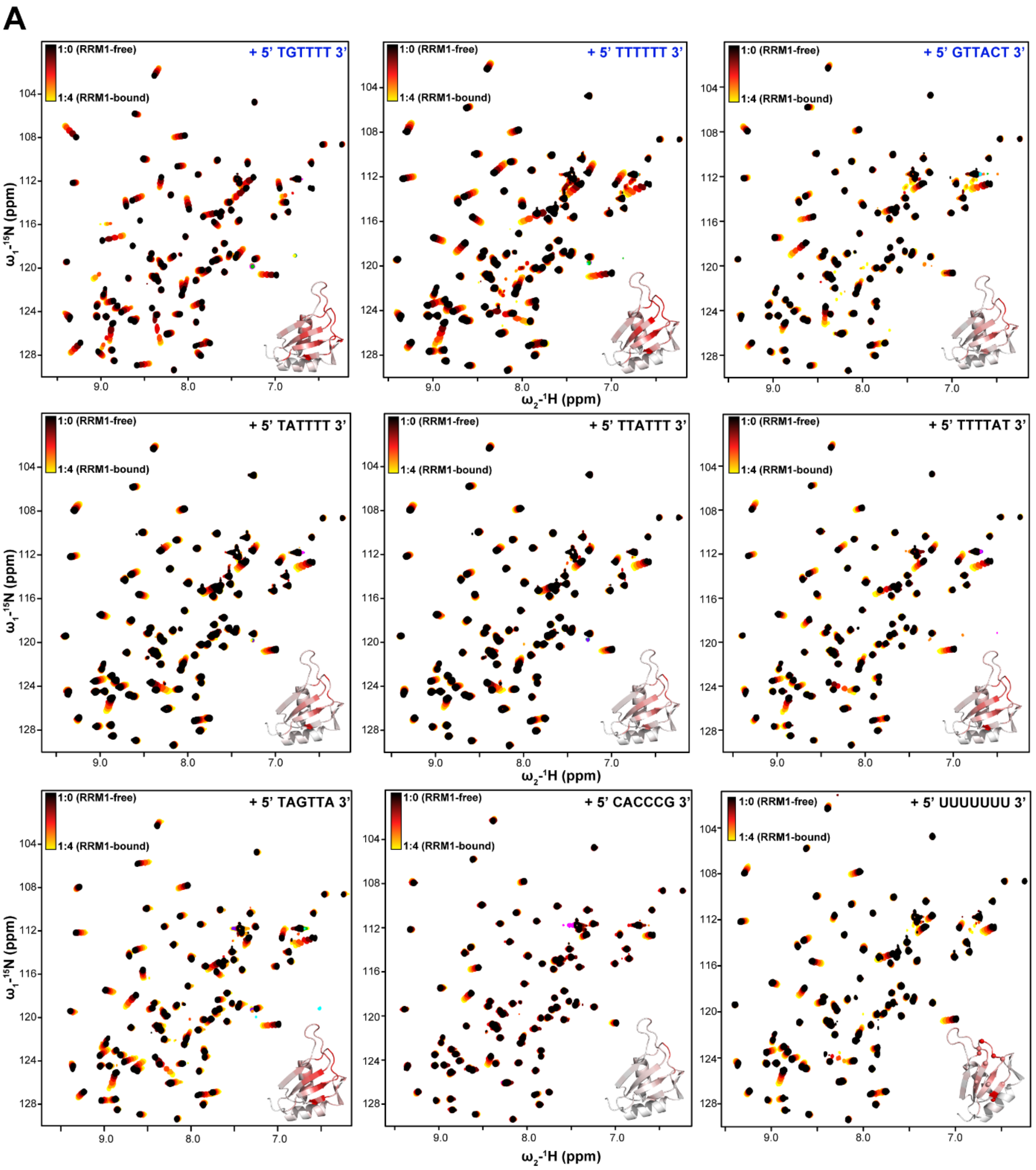
A
1:0 (RRM1-free)
1:4 (RRM1-bound)
+ 5' TGTTTT 3'
+ 5' TTTTTT 3'
+ 5' GTTACT 3'
+ 5' TATTTT 3'
+ 5' TTATTT 3'
+ 5' TTTTAT 3'
+ 5' TAGTTA 3'
+ 5' CACCCG 3'
+ 5' UUUUUUU 3'
ω1-15N (ppm)
ω2-1H (ppm)
104
108
112
116
120
124
128
9.0
8.0
7.0

**Figure S4B**

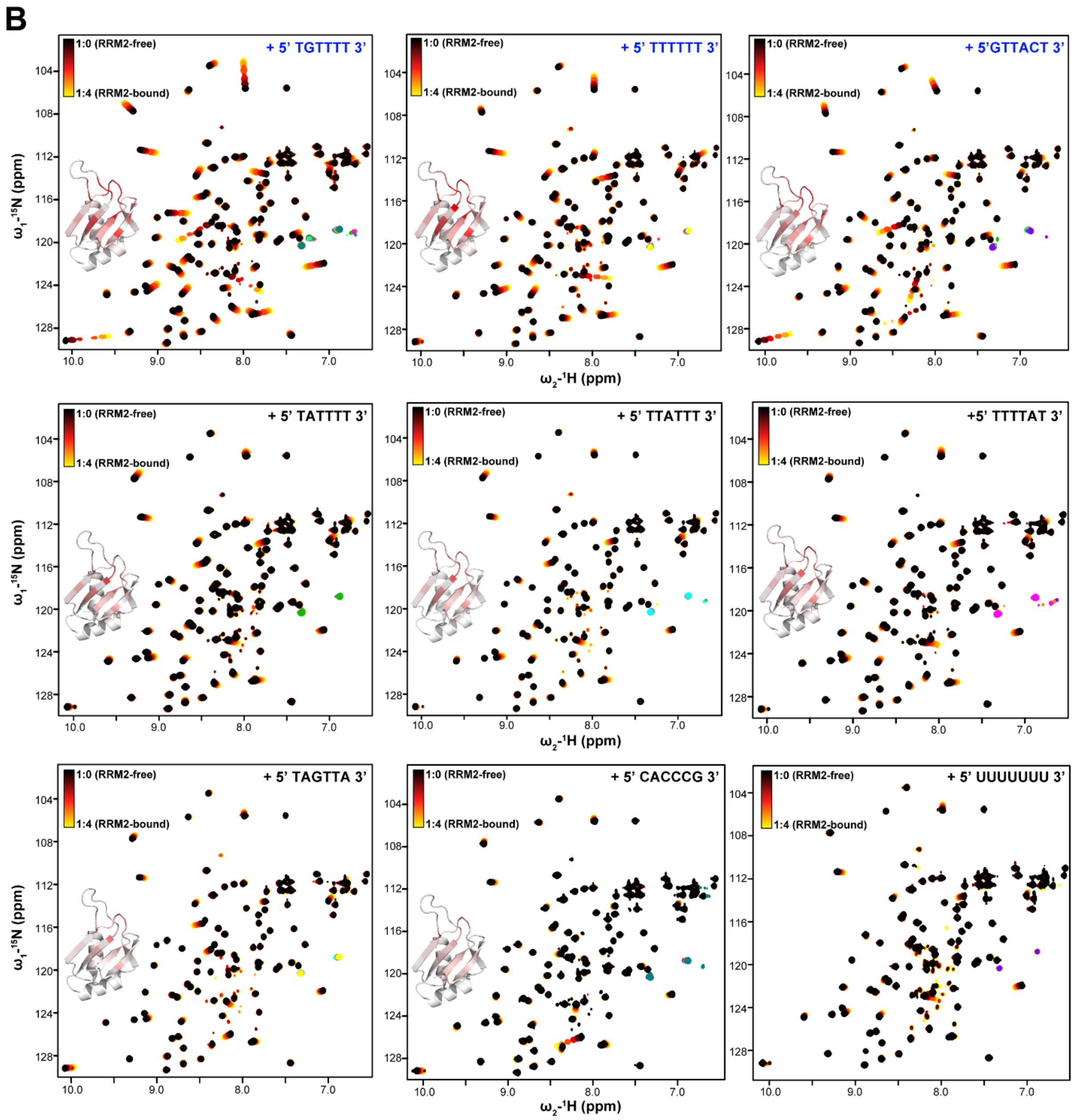

B
1:0 (RRM2-free)
1:4 (RRM2-bound)
+ 5' TGTTTT 3'
+ 5' TTTTTT 3'
+ 5'GTTACT 3'
+ 5' TATTTT 3'
+ 5' TTATTT 3'
+5' TTTTAT 3'
+ 5' TAGTTA 3'
+ 5' CACCCG 3'
+ 5' UUUUUUU 3'
ω1-15N (ppm)
ω2-1H (ppm)

**Figure S4C,D**

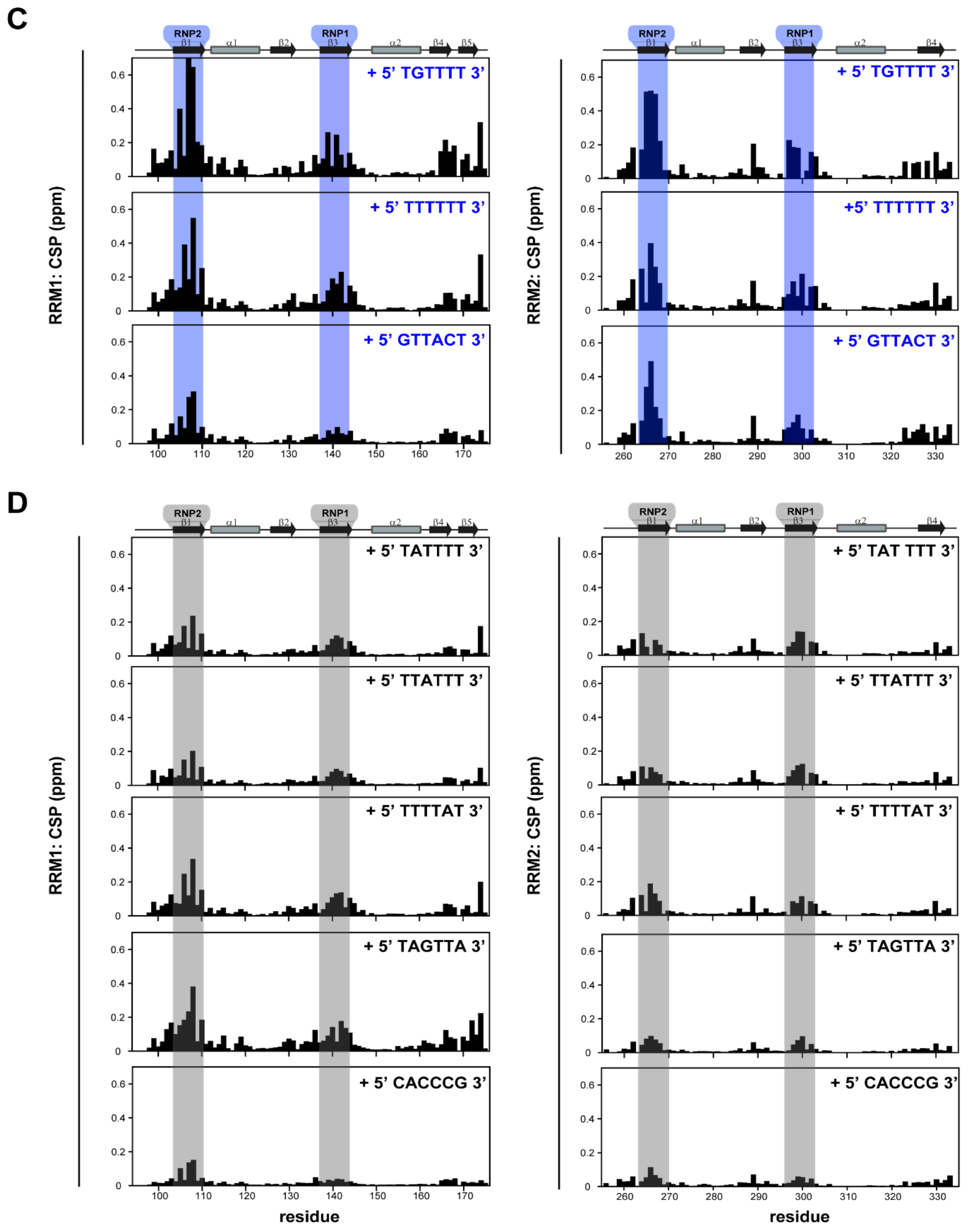


**Figure S4: NMR-based binding analysis of RRM1 and RRM2 of CP29a with various "T"-rich short oligonucleotide motifs using solution NMR.** Overlays of $^{1}H$-$^{15}N$ HSQC spectra of (A) RRM1 and (B) RRM2 of CP29a are shown with an increasing concentration of 6mer "TGTTTT", "TTTTTT", "GTTACT", "TATTTT", "TTATTT", "TTTTAT", "TAGTTA" and "CACCCG" single-stranded DNA oligonucleotides derived from CLIP-based analysis respectively. The change in resonances upon oligo. binding are colored with black (free-form) to red (intermediate bound-form) to yellow (oligo bound-form) gradient. The change in chemical shifts upon oligo. bindings are also mapped on the structural model of RRM1 of CP29a with gray (no-binding) to red (binding) color gradient for respective spectra. (C) Chemical shift perturbation (CSP) plots for RRM1 (left panel) and RRM2 (right panel) upon binding with 4-fold excess of oligos are shown. RRM1 and RRM2 domains show strong shifts for "TGTTTT", "TTTTTT" and "GTTACT" motifs. Binding sites on RNP1 and RNP2 regions are highlighted with blue. Secondary structural elements are shown on the top of the plot. (D) Similar CSP plots and analysis are derived with other variants of oligos.

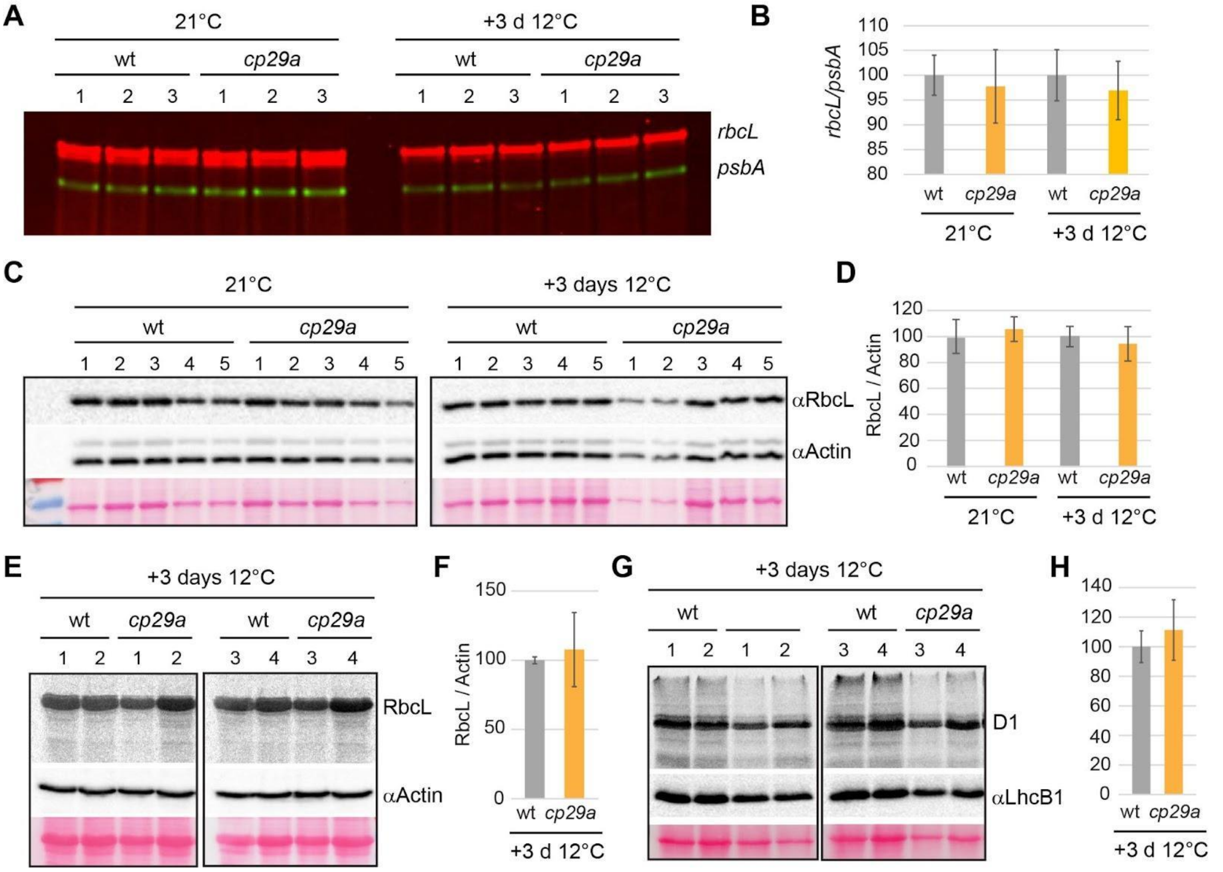


**Figure S5: Analysis of RbcL expression in *Arabidopsis cp29a* mutants.** (A) Two µg total leaf RNA from wt and *cp29a* mutants grown under standard conditions for two weeks or grown for the same conditions but then exposed for three days to 12°C (+3 d 12°C) were analyzed by RNA gel blot hybridization. Blots were probed simultaneously for the *psbA* and *rbcL* mRNAs using different fluorescence labels for probe preparation. Note that there are two isoforms of the *rcbL* mRNA accumulating under normal conditions, while the smaller isoform is strongly reduced in both wt and the *cp29a* mutant in the cold. Numbers indicate RNA preparations from independent plants. (B) The ratio of the *rbcL* over *psbA* mRNA signal was calculated for wt and mutants under the two conditions tested. (C) Immunoblot analysis of total protein preparations from wt and *cp29a* mutants grown under the two conditions described in (A). Antibodies against RbcL and actin were used consecutively on the same blots. (D) RbcL signals were normalized to actin signals. (E) In vivo pulse-labeling of leaf proteins from plants grown under short-term cold acclimation conditions. Leaf proteins were radiolabeled for 20 min with [35S] methionine. Total proteins were fractionated into soluble and insoluble proteins. Here soluble proteins were separated by SDS-PAGE and blotted onto a membrane prior to detection of radio-signals. The most prominent band is RbcL. The blot was afterwards probed with an antiserum directed against actin. (F) The RbcL signal was normalized using the nucleus-encoded actin protein. (G) For the detection of the membrane protein D1, the insoluble proteins of the preparation shown in (E) were analyzed by pulse-labeling. An antiserum against nucleus-encoded LhcB1 was used as a control for loading. (H) The D1 signal was normalized using the LhcB1 protein.

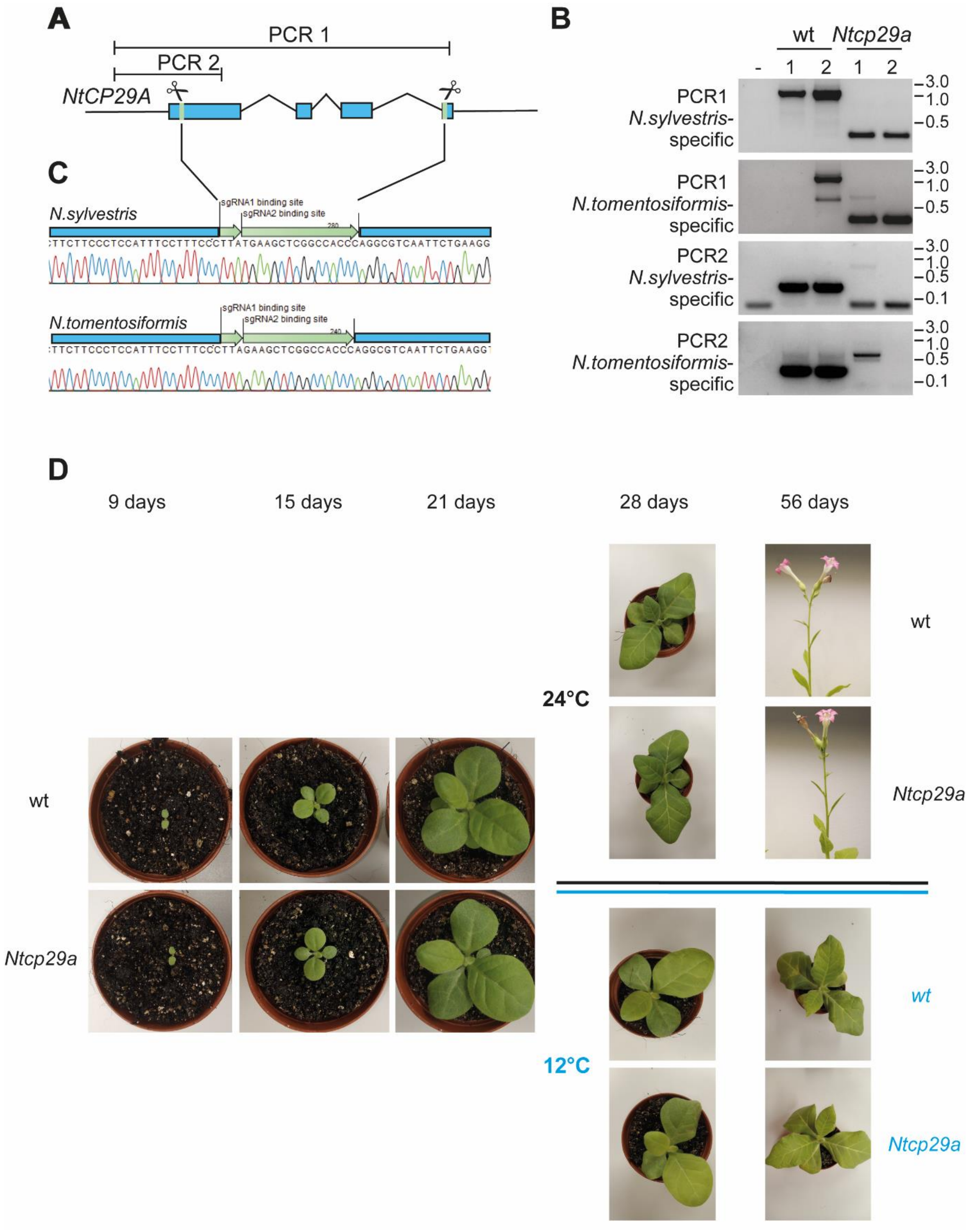


**Figure S6: Targeted deletion of NtCP29A in tobacco using CRISPR/Cas9.** (A) Schematic representation of the gene structure of *NtCP29A* and position of target sites of the guide RNAs (indicated by green lines and scissors) located on the first and fourth exons, respectively. (B) PCR analysis of leaf tissue of two independent transgenic lines showed the deletion events (i.e., bands shifted <500 bp) in *Ntcp29a* plants. Primer positions chosen allowed differentiation between the *N. sylvestris* and *N. tomentosiformis* alleles. In wt, all alleles were detected with the expected size for the amplification products, while the signals are absent in the two individual plants from the CRISPR/Cas9 mutant line. The minus sign denotes a control PCR reaction with water versus any DNA sample added. (C) Sequence analysis of the *Ntcp29a* deletion line. Sequences of the single guide RNA (sgRNA) target sites left after mutagenesis are indicated by green arrows. (D) Phenotype of wt and *Ntcp29a* mutants at normal growth temperatures and after shift to 12°C.

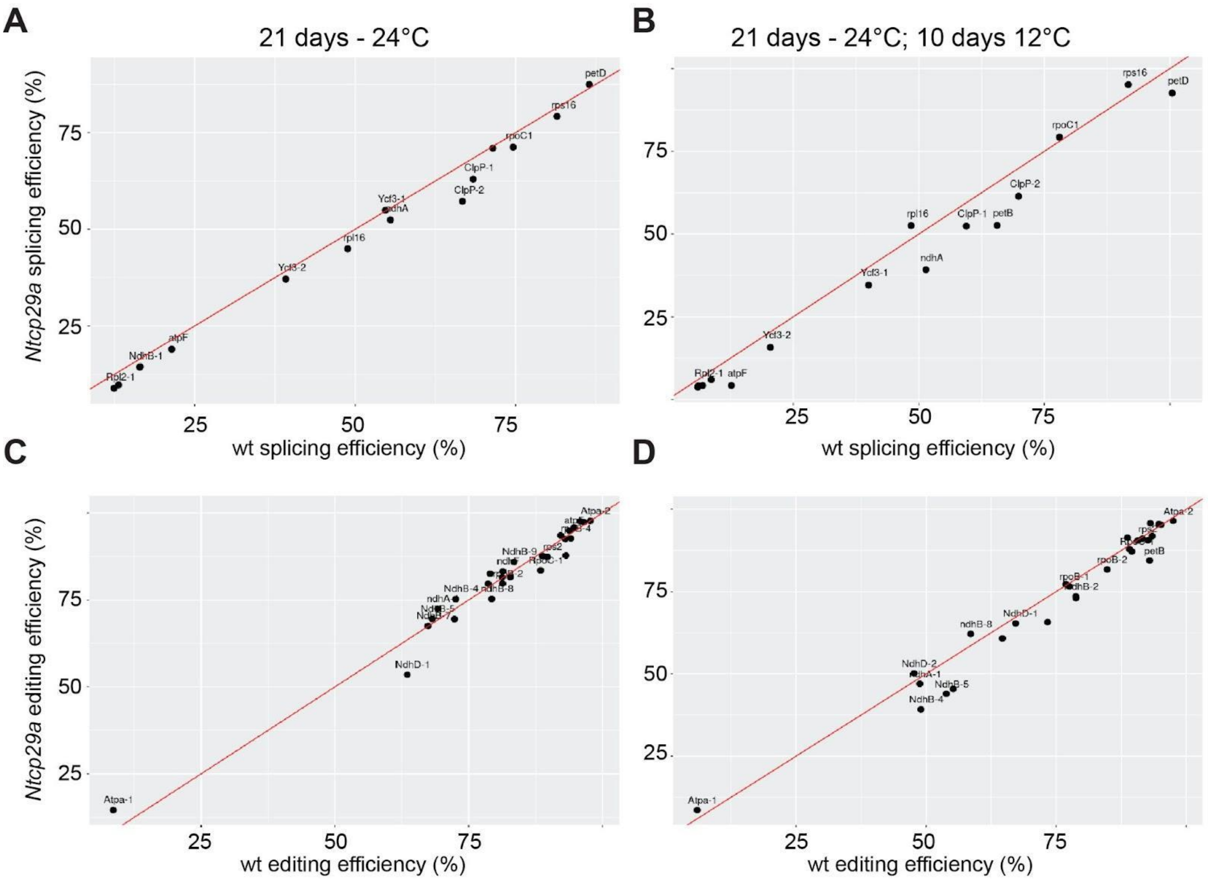


**Figure S7: Analysis of chloroplast RNA splicing and RNA editing in cold-treated *Ntcp29a* mutants.** (A) Analysis of splicing efficiency of chloroplast introns in wt and *Ntcp29a* mutant plants grown under standard conditions by RNA-seq. Splicing efficiency was calculated as the ratio of reads spanning exon-exon junctions versus reads spanning intron-exon boundaries (28). Using the Fisher's exact test in combination with Benjamini-Hochberg correction, we could not identify significant changes for any intron. (B) same analysis as in A, but based on samples grown for 21 days at 24°C and subsequently for 10 days at 12°C. (C) Analysis of RNA editing efficiency of chloroplast introns in wt and *Ntcp29a* mutant plants grown under standard conditions by RNA-seq. Editing efficiency was calculated as the ratio of edited versus unedited reads for known tobacco editing sites (28, 30). Fisher's exact test in combination with Benjamini-Hochberg correction did not identify significant changes for any editing site.

**Table S1: Top eCLIP peaks for AtCP29A.**

| Gene | Start[1] | Stop[1] | Length | $log_2$ (IP/input)[2] | $log_{10}$ p value[3] | Strand[1] |
|---|---|---|---|---|---|---|
| *rbcL 5'UTR-1* | 54945 | 54951 | 7 | 2.66 | 58.20 | + |
| *rbcL 5'UTR-2* | 54880 | 54896 | 72 | 3.51 | 57.61 | + |
| *trnH* | 19 | 46 | 28 | 3.19 | 23.06 | - |
| *psbL* | 63831 | 63881 | 51 | 3.85 | 17.13 | - |
| *psbB-1* | 73108 | 73144 | 37 | 2.70 | 14.97 | + |
| *psbT* | 74117 | 74161 | 45 | 4.20 | 12.68 | + |
| *atpF* | 11525 | 11561 | 37 | 4.11 | 11.74 | - |
| *psbB-2* | 73607 | 73616 | 10 | 3.29 | 11.56 | + |
| *psbD* | 33181 | 33188 | 8 | 2.66 | 11.51 | + |
| *petB* | 76214 | 76242 | 29 | 2.40 | 11.26 | + |

[1] GenBank acc. no. NC_000932
[3] The number of eCLIP reads overlapping CLIPper-identified peaks and the number overlapping the identical genomic region in the paired size-matched Input sample were counted and used to calculate fold enrichment (normalized by total usable read counts in each data set).
[2] enrichment p-value of reads in the eCLIP precipitated versus reads in the size-matched input calculated by Yates' Chi-Square test.

### Supplementary Materials and Methods

### Section I

#### eCLIP analysis

In this experiment, approximately $4\times10^9$ isolated chloroplasts were resuspended in RB-buffer (0.3 M sorbitol, 20 mM tricine-KOH (pH 8.4), 2.5 mM EDTA, 5 mM MgCl2) and placed onto a glass petri dish. The chloroplasts were then exposed to 500 $mJcm^{-2}$ of UV light at a wavelength of 254 nm while being kept on ice. After UV exposure, the chloroplasts were centrifuged at 500 × g for 5 minutes at 4°C, forming a pellet. The supernatant was discarded, and the resulting pellet was flash-frozen in liquid nitrogen and stored at -80°C. For further processing, the chloroplast pellets were thawed on ice and resuspended in 2 ml of CLIP lysis-buffer each (50 mM Tris-HCl (pH 7.4), 100 mM NaCl, 0.5% Nonidet P-40, 0.5% sodium deoxycholate, 1xcOmpleteTM EDTA-free Protease Inhibitor Cocktail; Roche). The resulting lysates were then centrifuged at 20,000 × g at 4°C for 30 minutes. Specific antibodies targeting the RNA-binding protein (RBP) of interest (10 µl for both anti-CP29A and anti-CP33B) were attached to 50 µl of Dynabeads Protein G (Invitrogen) and then resuspended in 500 µl of CO-IP buffer (150 mM NaCl, 20 mM Tris-HCl pH (7.5), 2 mM MgCl2, 5 µg/mL aprotinin, 0.5% Nonidet P-40). The supernatant from the chloroplast lysate pellet was transferred to a new tube, combined with 500 µl of the prepared bead suspension, and 4 units of Turbo DNase (Invitrogen). This mixture was incubated for 1 hour at 4°C with rotation. After incubation, 5% of the bead-lysate mixture was set aside for preparing size-matched input libraries and for western blot analysis of the immunoprecipitation. Finally, the supernatant was removed from the post-immunoprecipitation mixture and retained for western blot analysis.
The library preparation protocol for this study closely followed the established eCLIP protocol (12). Initially, the beads were washed twice with high-salt CLIP washing buffer (1000 mM NaCl, 20 mM Tris-HCl (pH 7.4), 1 mM EDTA, 0.5% Nonidet P-40), once with CLIP washing buffer, and once with FastAP buffer (10 mM Tris-HCl (pH 7.5), 5 mM MgCl2, 100 KCl, 0.02% Triton X-100). On-bead RNA dephosphorylation was performed by incubating with FastAP at 37°C for 15 minutes, followed by a 20-minute incubation with T4 polynucleotide kinase at the same temperature. Turbo DNase was added to both dephosphorylation steps. Subsequently, the

beads underwent two washes in CLIP washing buffer (20 mM Tris-HCl (pH 7.4), 10 mM MgCl2, 0.5% Nonidet P-40) and one in T4 RNA ligase 1 buffer (50 mM Tris-HCl (pH 7.5), 10 mM MgCl2; omitting DTT). A color-balanced pair of two 3'-CLIP-RNA adapters was ligated on-bead to the RNAs in each sample using T4 RNA ligase 1 (New England Biolabs), with the ligation reaction incubated for 75 minutes at 21°C. Post-ligation, the beads were washed three times in CLIP washing buffer. Both the crosslinked, adapter-ligated protein-RNA complexes and the previously set aside input samples were size-separated via polyacrylamide gel electrophoresis and transferred to nitrocellulose membranes. The RNA, positioned 75 kDa above the RBP of interest, was released from the nitrocellulose membrane using Proteinase K (New England Biolabs) digestion. The released RNA of the size-matched input samples was dephosphorylated and adapter-ligated, similar to the IP samples, but using a single 3'-input-RNA adapter. Both the IP and size-matched input samples' adapter-ligated RNA was reverse transcribed using SuperScript II Reverse Transcriptase (Invitrogen) at 42°C for 45 minutes. Following ExoSAP-IT (Applied Biosystems) treatment and chemical hydrolysis of residual RNA, cDNA was recovered using MyONE Silane beads (ThermoFisher Scientific). A 5'-DNA adapter, including unique molecular identifiers, was ligated to all cDNA samples using T4 RNA ligase 1 (New England Biolabs). This ligated cDNA was purified again using MyONE Silane beads (ThermoFisher Scientific). Quantification of the cDNA samples was done via qPCR, followed by PCR amplification using Q5 High-Fidelity DNA Polymerase (New England Biolabs) and indexed primers for Illumina sequencing. The resulting libraries were purified using the GeneJET PCR Purification Kit (ThermoFisher Scientific) and separated on 6% polyacrylamide gels. Library fragments ranging from 170 bp to 350 bp were extracted and subjected to Illumina sequencing.

**Computational analysis of eCLIP data**

The analysis of the eCLIP libraries and corresponding size-matched input controls was conducted using the CWL-based and dockerized eCLIP pipeline version 0.3.99 (available at github.com/YeoLab/eclip/releases/tag/v0.3.99). The general methodology followed the approach described previously (12), with some minor modifications. A custom docker container was created, incorporating an annotation of the *Arabidopsis thaliana* chloroplast chromosome for use with the CLIPPER tool. This custom annotation was derived from the AtRTD2 annotation, enhanced by extending all chloroplast coding sequences by 100 base pairs at both the 5'- and 3'-UTR ends. The criteria for identifying significant peaks in the analysis were stringent: a minimum of 4-fold enrichment of IP over the input control was required, along with an adjusted p-value of ≤ 0.001. Additionally, peaks were only considered significant if they were identified as such in both biological replicates.

**RBNS analysis**

We slightly modified the RNA Bind-n-Seq protocol (17). AtCP29A and AtCP33B were tagged with N-terminal streptavidin-binding protein (SBP) and produced using the pGEX system. Their quality and quantity were verified by SDS-PAGE and a protein standard dilution series. The *in vitro* transcribed RNA pool (0.5 µM) was then mixed with various concentrations of these RBPs (0, 10, 100 and 1000 µM) and incubated for 3 hours at 21°C. Magnetic Dynabeads MyOne Streptavidin C1 were then added, followed by another hour of incubation. The protein-RNA complexes were magnetically separated and washed twice. RNA was eluted in SDS-containing buffer at 70°C for 10 minutes and purified using the RNA Clean & Concentrator-5 kit. The RNA was reverse transcribed using ProtoScript II Reverse Transcriptase, and 0.5 pm of the RNA pool was also reverse transcribed. cDNA libraries were amplified with NEXTflex® Unique Dual Index Barcodes and Q5® High-Fidelity DNA Polymerase, then size-selected and purified via gel electrophoresis for sequencing on the Illumina NextSeq500 platform. Binding-buffer composition was 25 mM Tris-HCl (pH 7.5), 150 mM KCl, 3 mM MgCl2, 0.01% Tween, and 1 mM DTT, 1 mg/mL BSA; Washing-buffer had 25 mM Tris-HCl (pH 7.5), 150 mM KCl, 0.5 mM EDTA, 0.01% Tween, 60 µg/mL BSA; Elution-buffer contained 10 mM Tris-HCl (pH 7.0), 1 mM EDTA, 1% SDS.

Data analysis RNA Bind-N-Seq libraries was performed using a published computational workflow (18, bitbucket.org/pfreese/rbns_pipeline). The kmer enrichment analysis was performed for k=6. We also checked 6, 7, 8, and 9 mers. All resulted in very similar variations of the 6-mer motifs, i.e. the polyU one for CP29A and the more complex one for CP33B. We also checked for gapped motifs, which resulted in the same pattern.

### Section II

#### Nuclear Magnetic Resonance (NMR) spectroscopy

N-terminal $His^6$-tagged RRM1 (97-176 aa) and RRM2 (255-334 aa) constructs were cloned into the bacterial expression vector pETM-11 and over-expressed in *E. coli* BL21 (DE3) in M9 minimal media supplemented with $^{15}N$-labeled $NH_4Cl$. A similar protein purification protocol was followed as described previously (16). Briefly, bacterial cells were lysed by French press and proteins were purified by Ni-NTA-based affinity chromatography followed by TEV cleavage, ion-exchange and size-exclusion chromatography. The final NMR buffer (20 mM sodium phosphate, pH 6.8, 50 mM NaCl, 1 mM DTT) was used for the protein samples.

All NMR measurements were carried out on Bruker NMR spectrometers with a proton Larmor frequency of 500 and 600 MHz equipped with cryogenic or room temperature $^1H$, $^{13}C$ and $^{15}N$ triple resonance probes. Shigemi tube was used for the sample measurement and 5% $D_2O$ was added to the samples to lock the external magnetic field. $^1H$, $^{15}N$ HSQC spectra were acquired at 298K temperature with 120 ms and 70 ms acquisition time in direct and indirect dimensions, respectively. Spectra were processed with Bruker Topspin 3.5pl6 software package using a shifted sine-bell window function and zero-filling before Fourier transformation. Proton chemical shifts were referenced against sodium 2,2-dimethyl-2-silapentane 5-sulfonate (DSS). All spectra were analyzed by the CCPN (v2.5) software package (50).

Protein backbone chemical shifts for RRM1 and RRM2 constructs were obtained from the BMRB (Biological Magnetic Resonance Data Bank) accession ID 52022 and 52025. For NMR titration experiments, series of $^1H$-$^{15}N$ HSQC experiments were measured using $^{15}N$-labeled 50 µM protein with stepwise increasing concentration of short RNA (Dharmacon, USA) or single-stranded DNA (Eurofins Genomics, Germany) oligonucleotides. Spectra were analyzed by CCPN and chemical shift perturbations (CSPs) were calculated between oligo-free form and oligo-bound form (4-fold molar excess) of RRM using the equation: $\Delta\Delta\delta = [\,(\Delta\delta^1H)^2 + (\Delta\delta^{15}N/5)^2\,]^{1/2}$. The derived CSPs were mapped on the RRM structural model and further analyzed by Schrodinger's PyMol tool. Cumulative CSPs of each dataset were further calculated by average maximum shifts observed on RNP2 regions of the RRM and analyzed. NMR-based dissociation constants ($K_D$) were derived by fitting to the equation:

$\Delta\delta_{obs} = \Delta\delta_{max}\ \{\ ([P]_t + [L]_t + K_D) - [\ ([P]_t + [L]_t + K_D)^2 - 4[P]_t\ [L]_t]^{½}\ \}\ /\ 2[P]_t$, where, $\Delta\delta_{obs}$ is the observed chemical shift difference in each titration point relative to the free state, $\Delta\delta_{max}$ is the maximum shift change at 4-fold excess of oligo, $[P]_t$ and $[L]_t$ are the total protein and oligo concentrations, respectively, and $K_D$ is the dissociation constant (51).

### Section III

#### Vector construction

#### CRISPR/Cas vector

Vector constructs were assembled as described (52).

#### Complementation constructs

The genomic DNA regions of NtsCP29A and NttCP29A including the UTRs and the promoter regions were amplified with gene-specific primers. The PCR products were cloned together with a spacer region into the pB7WG vector (53), which was then used in the tobacco transformation.

#### Tobacco transformation

Wild-type tobacco plants (Petit Havana) were transformed using the leaf disc transformation method.

### Section IV

#### Plant Material and Growth

Arabidopsis *cp29a-6* mutants were described previously (3). Arabidopsis seeds were grown at 21°C with a light intensity of 120 µmol $m^{-2}s^{-1}$. For cold treatment, plants were grown for 14 days at 21°C and then transferred to 8°C for 10 days. Tobacco plants were grown at 24°C with a light intensity of 300 µmol $m^{-2}s^{-1}$. For cold treatment, plants were grown for 21 days at 24°C and then transferred to 12°C for 7 days.

#### Chlorophyll fluorescence analysis

Chlorophyll a fluorescence *in vivo* was measured using the Imaging PAM chlorophyll fluorimeter (M-Series; Walz, Effeltrich, Germany). Measurements were performed following standard protocols (49).

#### RNA gel blot hybridization

Small RNA gel blot analysis was performed as described (21)) using an oligonucleotide probe with the sequence GCAATAAAACAAAACAACAAGGTCTACTCGACA. Standard RNA gel blot hybridization was carried out as described (3). RNA probes were prepared by *in vitro* transcription in the presence of 5-azido-C3-UTP nucleotides (Jena Biosciences) with final linking to sulpho-cyanine cy5.5 or cy7.5 dye (Lumiprobe) according to the manufacturer's protocol. Primer used for amplifying the probes are as following: CK_rbcL_for (GCAGCATTCCGAGTAACTCC) and CK_rbcL_T7 (GTAATACGACTCACTATAGGGCCACGTAGACATTCATAAACTGC) resulting in a fragment of 473bp covering the N terminal part of *rbcL* mRNA. The *psbA* probe was generated with the use of psbA.T7 primer (GTAATCGACTCACTATAGGGATTCCTAGAGGCATACCATCAG) and psbAfw primer (GAAAGCGAAAGCCTATGGGG) resulting in a PCR fragment of 503bp.

#### Small RNA sequencing

Small RNA sequencing and bioinformatic analysis were done as described earlier (54).

#### RNA Seq analysis

RNA Seq data were processed using the nf-core/rnaseq (https://github.com/nf-core/rnaseq/) pipeline. Reads were mapped to the tobacco chloroplast (*Nicotiana tabacum* plastid, NC_001879.2). FeatureCounts (Rsubread, v2.0.6) was used for counting. rRNAs and tRNAs were removed from further analysis. Differentially expression analysis was done with DESeq2 (29). Splicing and editing efficiencies were calculated using the chloroseq pipeline (28).

#### Immunoblot analysis

Total protein preparations were loaded based on equal fresh weight. The proteins were separated by SDS-PAGE and transferred to a PVDF membrane. Protein integrity and loading were tested by Ponceau S staining. Hybridization with the primary antibody was carried out overnight at 4°C and with the second antibody for 1 hour at room temperature. Stripping of antibodies was done with 1.5% glycine, 0.1% SDS, 1 & Tween 20, pH = 2.2.

#### *In Vivo* Pulse-Chase Labeling of Chloroplast-Encoded Proteins

Young leaves from 21 day old Arabidopsis plants were used for $^{35}$S-methionine-labeling of proteins as previously described (55).
